# Estrogen receptor alpha controls gene expression via translational offsetting

**DOI:** 10.1101/507574

**Authors:** Julie Lorent, Richard J. Rebello, Vincent van Hoef, Mitchell G. Lawrence, Krzysztof J. Szkop, Eric Kusnadi, Baila Samreen, Preetika Balanathan, Karin Scharmann, Itsuhiro Takizawa, Sebastian A. Leidel, Gail P. Risbridger, Ivan Topisirovic, Ola Larsson, Luc Furic

**Author notes:** Equally contributing first authors (Julie Lorent, Richard J. Rebello, Vincent van Hoef). Deceased author (Itsuhiro Takizawa).

## Abstract

Estrogen receptor alpha (ERα) activity is associated with increased cancer cell proliferation. Studies aiming to understand the impact of ERα on cancer-associated phenotypes have largely been limited to its transcriptional activity. Herein, we demonstrate that ERα coordinates its transcriptional output with selective modulation of mRNA translation. Importantly, translational perturbations caused by depletion of ERα largely manifest as “translational offsetting” of the transcriptome, whereby amounts of translated mRNA and protein levels are maintained constant despite changes in mRNA abundance. Transcripts whose levels, but not polysome-association, are reduced following ERα depletion lack features which limit translational efficiency including structured 5’UTRs and miRNA target sites. In contrast, mRNAs induced upon ERα depletion whose polysome-association remains unaltered are enriched in codons requiring U34-modified tRNAs for efficient decoding. Consistently, ERα regulates levels of U34-modification enzymes, whereas altered expression of U34-modification enzymes disrupts ERα dependent translational offsetting. Altogether, we unravel a hitherto unprecedented mechanism of ERα-dependent orchestration of transcriptional and translational programs, and highlight that translational offsetting may be a pervasive mechanism of proteome maintenance in hormone-dependent cancers.

## Introduction

Gene expression is regulated at multiple levels including transcription and mRNA transport, storage, stability and translation. These processes together with protein degradation govern proteome composition (Morris *et al*, 2010; Piccirillo *et al*, 2014; Bisogno & Keene, 2018). While mRNA levels are key determinants of the proteome under non-stressed growth conditions, the contribution of mRNA translation remains contentious (Schwanhäusser *et al*, 2011; Li *et al*, 2014, 2017). In contrast, it is well established that altered translational efficiencies reshape the proteome during various dynamic responses including cellular differentiation and endoplasmic reticulum stress (Kristensen *et al*, 2013; Baird *et al*, 2014; Liu *et al*, 2016; Guan *et al*, 2017).

Translation can be regulated globally, leading to altered translational efficiency of most cellular mRNAs, or selectively by modulating translation of limited subsets of mRNAs (Piccirillo *et al*, 2014). Most commonly, selective changes in translational efficiency are considered to allow modulation of protein levels in absence of corresponding changes in mRNA levels (Larsson *et al*, 2010). This is thought to be mediated by interactions between RNA-elements in untranslated regions (UTRs), RNA-binding proteins and translation initiation factors (Koromilas *et al*, 1992; Hershey *et al*, 2012; Hinnebusch *et al*, 2016; Masvidal *et al*, 2017). Accordingly, 5’UTR length, structure and presence of cis-acting elements such as upstream open reading frames (uORFs); and 3’UTR trans-acting factors including microRNAs (miRNAs) play a pivotal role in translational control (Gebauer *et al*, 2012; Larsson *et al*, 2013; Hinnebusch *et al*, 2016; Gandin *et al*, 2016b). Although the vast majority of known RNA elements implicated in modulation of mRNA translation reside within UTRs, it has been reported that nucleotide sequence and/or elements in the open reading frame may also regulate translation (López *et al*, 2015; Thandapani *et al*, 2015). Indeed, selective alterations of transfer RNAs (tRNAs) were recently described to affect translation of mRNAs with specific codon usage (Goodarzi *et al*, 2016). Moreover, recent reports highlight alterations of tRNA modifications as an important mechanism underlying selective translation (Chan *et al*, 2015; Delaunay *et al*, 2016; Rapino *et al*, 2017, 2018) which are thought to act by maintaining protein homeostasis or driving an adaptive proteome (Nedialkova & Leidel, 2015).

Estrogen receptor alpha (ERα) is a key steroid receptor and transcription factor which drives tumorigenesis in hormone-dependent cancers (Shanle & Xu, 2010). In prostate cancer, ERα expression is associated with increased cell proliferation. Moreover, ERα is overexpressed in genetically engineered mouse models of prostate cancer and high grade patient tumors (Chakravarty *et al*, 2014; Megas *et al*, 2015; Takizawa *et al*, 2015). In this context, the ERα transcriptional output is thought to direct a program distinct from the androgen receptor (AR) that may contribute to emergence of castrate-resistant prostate cancer and aggressive tumor subtypes (Setlur *et al*, 2008). Intriguingly, in addition to its role in regulating transcription, ERα may directly or indirectly influence PI3K/AKT/mTOR and MAPK pathway signaling, as shown in several tissues (Levin, 2009) including the prostate (Takizawa *et al*, 2015). Studies in multiple cellular models have revealed that fluctuations in mTOR activity predominantly affect translation of mRNA subsets defined by long and highly structured 5’UTRs, extremely short 5’UTRs or 5’UTRs harboring a 5’terminal oligopyrimidine tract (TOP) (Patursky-Polischuk *et al*, 2009; Hsieh *et al*, 2012; Larsson *et al*, 2012; Meyuhas & Kahan, 2015; Gandin *et al*, 2016b; Masvidal *et al*, 2017). Moreover, MAPK signaling induces phosphorylation of eIF4E which also selectively modulates mRNA translation (Furic *et al*, 2010; Robichaud *et al*, 2015). Therefore, we investigated whether ERα, in addition to its well-established role in transcription, also modulates translation.

## Results

### Changes in steady-state mRNA levels upon depletion of ERα are largely offset at the level of translation

To study the impact of ERα on regulation of gene expression in prostate cancer, we used the BM67 cell line derived from the PTEN null mouse model of prostate cancer (Takizawa *et al*, 2015). BM67 cells express relatively high level of ERα, which was silenced using an shRNA to generate shERα BM67 cells (**Fig. 1A**). To assess effects of ERα depletion on mRNA abundance and translation, we used polysome-profiling quantified by DNA-microarrays (**Supplementary Fig. S1A-B**). Polysome-profiling generates parallel data on efficiently translated (i.e. those associated with >3 ribosomes) and total mRNA (**Fig. 1A**) (Gandin *et al*, 2016b; Masvidal *et al*, 2017). Changes in polysome-associated mRNA can result either from congruent changes in cytosolic mRNA or from changes in translational efficiencies without corresponding fluctuations in mRNA levels. The former is the outcome of regulated transcription and/or mRNA stability while the latter reflects *bona fide* changes in translational efficiency (Larsson *et al*, 2010). The anota2seq algorithm identifies changes in polysome-associated and cytosolic mRNA; and computes alterations in translational efficiency (Larsson *et al*, 2010; Oertlin *et al*, 2018). In line with the role of ERα as a transcription factor, wide-spread changes in total mRNA levels were observed between control and shERα BM67 cells as evidenced by a strong enrichment of transcripts with low p-values (**Fig. 1B**; **Supplementary Fig. S1C**). Substantially fewer mRNAs displayed ERα-associated changes in polysome-association as indicated by a smaller enrichment of transcripts with low p-values (**Fig. 1B**; **Supplementary Fig. S1C**). Strikingly, when comparing fold changes for cytosolic and polysome-associated mRNAs in shERα vs. control BM67 cells, we identified a population of genes showing changes in cytosolic mRNA without corresponding changes in their association with polysomes (**Fig. 1C**). This suggests that shERα-dependent alterations in mRNA levels may be buffered at the level of translation such that the amount of mRNA associated with polysomes is unaltered despite changes in corresponding mRNA levels. Translational buffering has been described in the context of transcript-dosage compensation where it acts to maintain protein levels similar between different bacterial (Lalanne *et al*, 2018) and yeast species (Artieri & Fraser, 2014; McManus *et al*, 2014) and human individuals (Cenik *et al*, 2015). It has also been reported that gene dosage effects caused by aneuploidy may be compensated at the level of translation in a cell type-specific context (Zhang & Presgraves, 2017), and that translational buffering may reduce transcriptional “noise” caused by acute stimulation of cells with growth factors (Tebaldi *et al*, 2012). In contrast to these modes of regulation, which appear to chiefly reduce “noise” in the proteome composition, transcriptional defects caused by ERα-depletion appear to induce a form of translational buffering which can be activated to sustain an adaptive proteome and thus we refer to it as “translational offsetting”.

**Figure 1.**
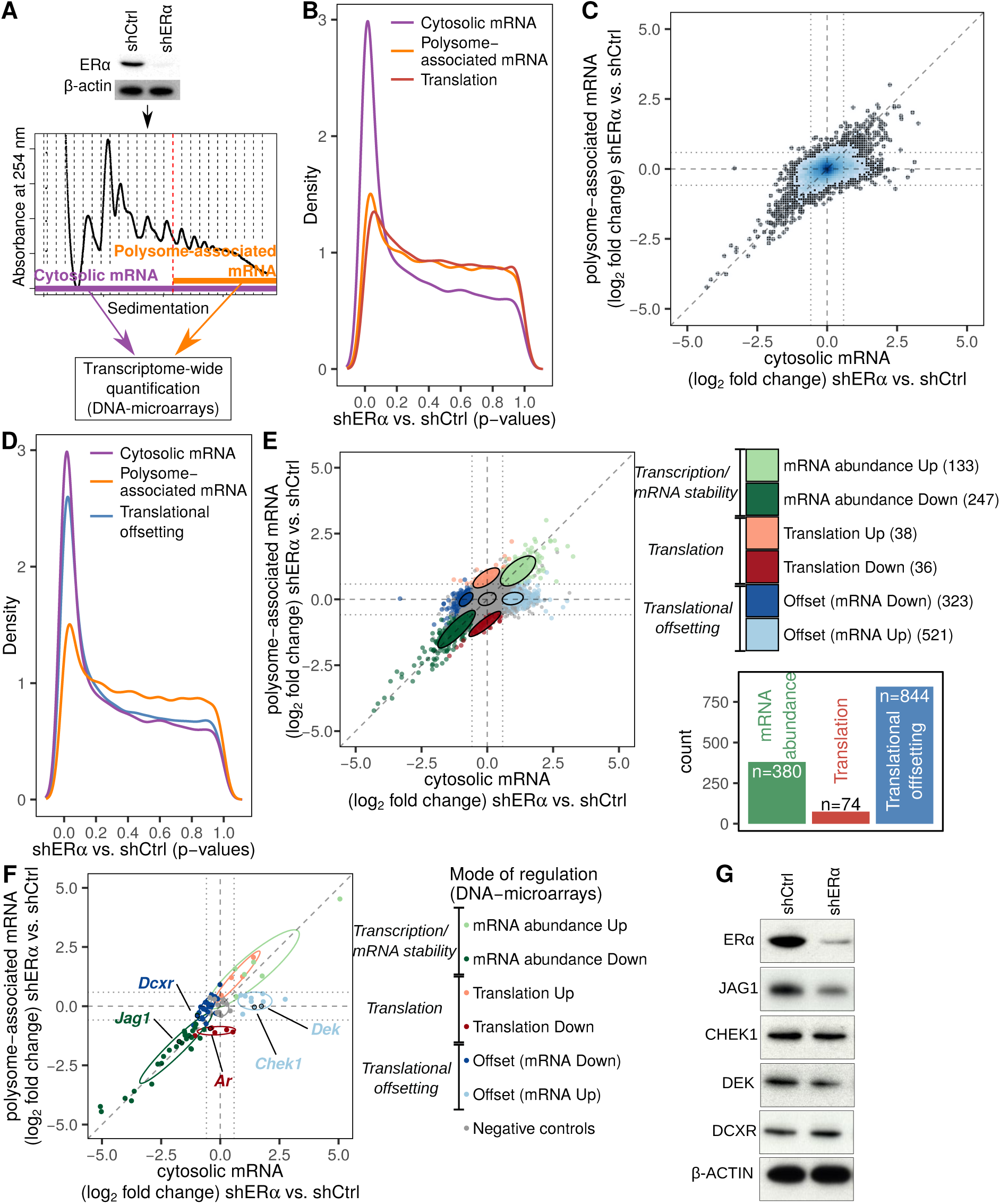
ERα-dependent alterations in steady-state mRNA levels are largely offset at the level of mRNA translation. **(A)** Expression of ERα in control (shCtrl) and knockdown (shERα) BM67 cells was determined by western blotting (β-actin served as a loading control). Gene expression was determined using polysome-profiling which quantifies both total mRNA and efficiently translated polysome-associated mRNA. **(B)** Densities of p-values for differential expression (shERα vs. shCtrl BM67 cells) using data from polysome-associated mRNA (orange), cytosolic mRNA (purple) or from analysis of changes in translational efficiency leading to altered protein levels (red). **(C)** Scatterplot of polysome-associated mRNA vs. cytosolic mRNA log2 fold-changes (shERα vs. shCtrl). Areas of the plot are colored according to the density of data points (genes; dark blue corresponds to areas with many genes [see methods]) **(D)** Same plot as in (B) but including a blue density of p-values from the analysis of translational offsetting. **(E)** A scatter plot similar to (C) but where genes are colored according to their mode of regulation derived from anota2seq analysis. A relaxed threshold (p<0.05) was used to identify a set of transcripts regulated via translation, which did not pass thresholds used for identification of changes in mRNA abundance or translational offsetting (FDR<0.1). Confidence ellipses (level 0.7) are overlaid for each mode of regulation. A bar graph indicates the number of mRNAs regulated via each mode. **(F)** Targets from each mode of regulation (cf. E) were selected for validation by Nanostring. Shown is a scatterplot of Nanostring quantification of polysome-associated mRNA vs. cytosolic mRNA log2 fold-change (shERα vs. shCtrl). Genes are colored according to their identified mode of regulation (i.e. from E). Confidence ellipses (based on Nanostring data, level 0.7) are overlaid for each mode of regulation. Selected genes from each mode of regulation are indicated. **(G)** Levels of indicated proteins from shERα and control BM67 cells were determined by western blotting. β-actin served as loading control.

We next implemented adjustments in anota2seq that allowed analysis of different forms of translational buffering including ERα-dependent translational offsetting (Oertlin *et al*, 2018). This led to identification of a large number of mRNAs which changed in abundance in shERα BM67 vs. control cells but were translationally offset as illustrated by an abundance of low p-values (**Fig. 1D**; **Supplementary Fig. S1D**). Anota2seq also allows to categorize transcripts in 3 modes of regulation: i) changes in mRNA abundance (congruent changes in total and polysome-associated mRNA), ii) changes in translation (changes in polysome-associated mRNA without corresponding changes in total mRNA) and translational offsetting (changes in cytosolic mRNA without corresponding changes in their polysome association) (**Fig. 1E; Supplementary Table S1**). Strikingly, as evidenced by the number of transcripts under each mode of regulation, translational offsetting was the predominant mode for regulation of gene expression following ERα depletion (**Fig. 1E**). Therefore, the ERα-dependent perturbations in mRNA levels appear to be largely offset at the level of translation.

### ERα-dependent translational offsetting opposes changes in protein levels despite alterations in corresponding mRNA levels

To validate observed ERα-dependent changes in gene expression, we selected candidate genes from the three modes of regulation, together with negative controls. Nanostring technology (Geiss *et al*, 2008) quantified 86 such mRNAs and confirmed all modes of ERα-dependent regulation of gene expression (**Fig. 1F; Supplementary Fig. S2A.** We next selected genes regulated at the level of translation (AR), mRNA abundance (JAG1) or translational offsetting (CHEK1, DEK and DCXR) and assessed corresponding protein abundance using Western blotting. AR and JAG1 were downregulated in shERα as compared to control BM67 cells, which corresponded to the observed decrease in their polysome association (**Fig. 1G**, **Supplementary Fig. S2BC**). In turn, levels of proteins encoded by translationally offset mRNAs remained comparable between ERα depleted and control BM67 cells (**Fig. 1G, Supplementary Fig. S2B-C**). Altogether, in addition to changes in total mRNA levels, ERα depletion leads to translational offsetting which opposes changes in protein levels despite alterations in mRNA abundance.

We next sought to establish functional relationships between the ERα-dependent genes governed by translational offsetting by performing gene set enrichment analyses using Gene Ontology (GO) annotations (Gene Ontology Consortium, 2015). No functions passed selected thresholds for enrichment among proteins encoded by mRNAs induced in shERα vs. control BM67 cells but translationally offset. In contrast, metabolism and mitochondria-related functions were enriched (False Discovery Rate (FDR)-adjusted p-value = 0.008 and 0.001, respectively) among proteins encoded by mRNAs whose levels were suppressed but translationally offset upon ERα depletion (**Fig. 2; Supplementary Table S2**). As expected, this largely overlapped with the enrichment of cellular functions among proteins encoded by mRNAs whose total mRNA was reduced upon ERα depletion (irrespective of whether this was paralleled by changes in their polysome-association or not; **Supplementary Fig. S3; Fig. 2**). In contrast, there was no significant enrichment in cellular functions among proteins encoded by mRNAs whose polysome-association was reduced upon ERα depletion. Therefore, subsets of mRNAs which are offset at the level of translation upon ERα depletion are functionally related and are implicated in essential cellular functions.

**Figure 2.**
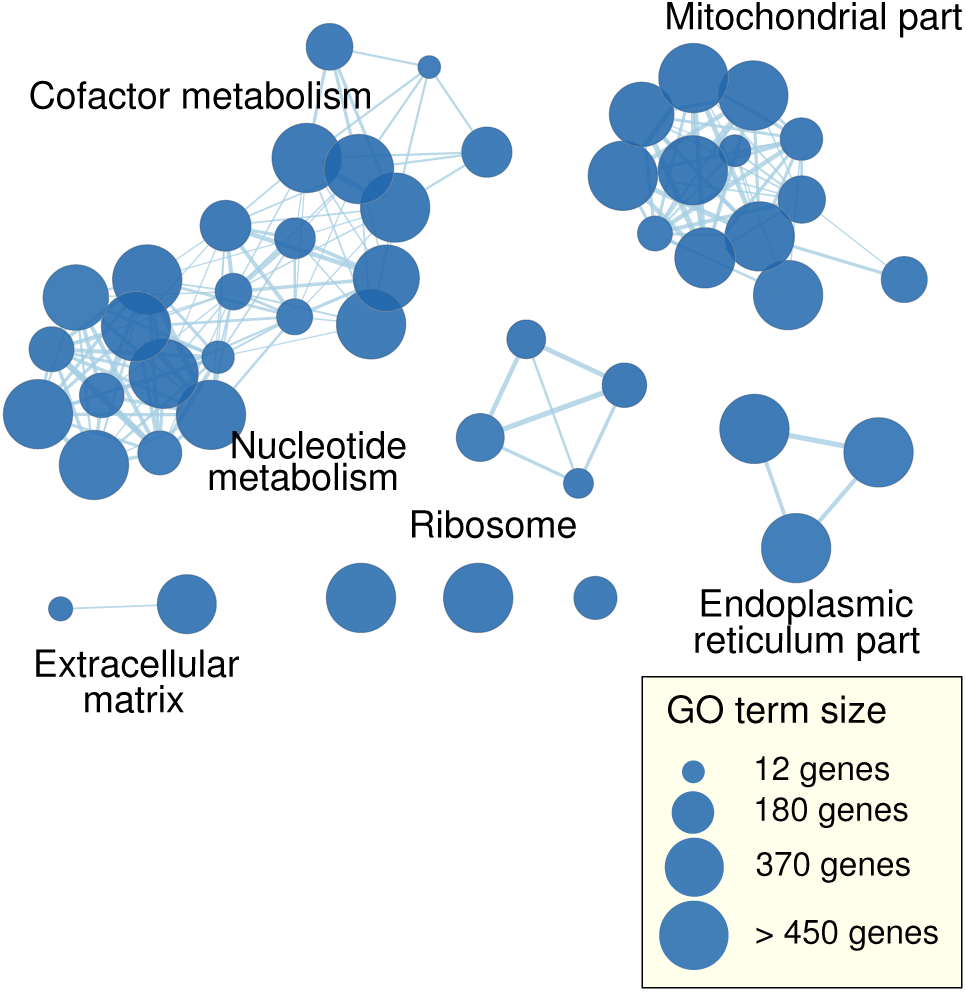
ERα depletion-associated translational offsetting modulates expression of genes belonging to distinct cellular functions. GO terms enriched among proteins encoded by mRNAs suppressed but offset upon ERα depletion with FDR < 0.15 are visualized as a network. Each node is a GO term connected to other GO terms whose enriched genes overlap. The size of each node reflects the number of genes associated to its corresponding term and the width of the edge illustrates the size of the overlap between two connected nodes.

### Short, unstructured 5’UTRs characterize mRNAs which are downregulated but translationally offset upon ERα depletion

We next sought to identify mRNA features which underpin ERα-dependent translational offsetting. To this end, we compared mRNAs which changed their level and polysome-association in ERα-depleted vs control cells vs. those that maintained their polysome occupancy despite changes in the abundance. We initially focused on 5’UTR features as they play key roles in translational control (Hinnebusch *et al*, 2016; Masvidal *et al*, 2017). To achieve this, we performed transcription start site profiling in shERα BM67 cells using nano-cap analysis of gene expression (nanoCAGE). At a sequencing depth close to saturation, approximately 10,000 5’UTRs were detected (**Fig. 3A**). Using these data, we contrasted genes whose mRNA abundance and polysome-association changed in parallel to those which were translationally offset for: 5’UTR length, GC content, free energy of folding and presence of uORFs in a strong Kozak context. Strikingly, transcripts whose levels were reduced upon ERα depletion but were translationally offset had a median 5’ UTR lengths ∼ 50% shorter and were less structured relative to downregulated but non-offset mRNAs (**Fig. 3B**). In contrast, transcripts induced upon ERα depletion but translationally offset exhibited comparable 5’UTR length and folding free energy but slightly lower GC content as compared to non-offset mRNAs (**Fig. 3B**). Furthermore, there were no differences in proportion of mRNA harboring uORFs between offset and non-offset transcripts (**Fig. 3B**). Collectively, these findings suggest that mRNAs whose level is suppressed upon ERα depletion but are offset at the level of translation contain shorter and less structured 5’UTRs.

**Figure 3.**
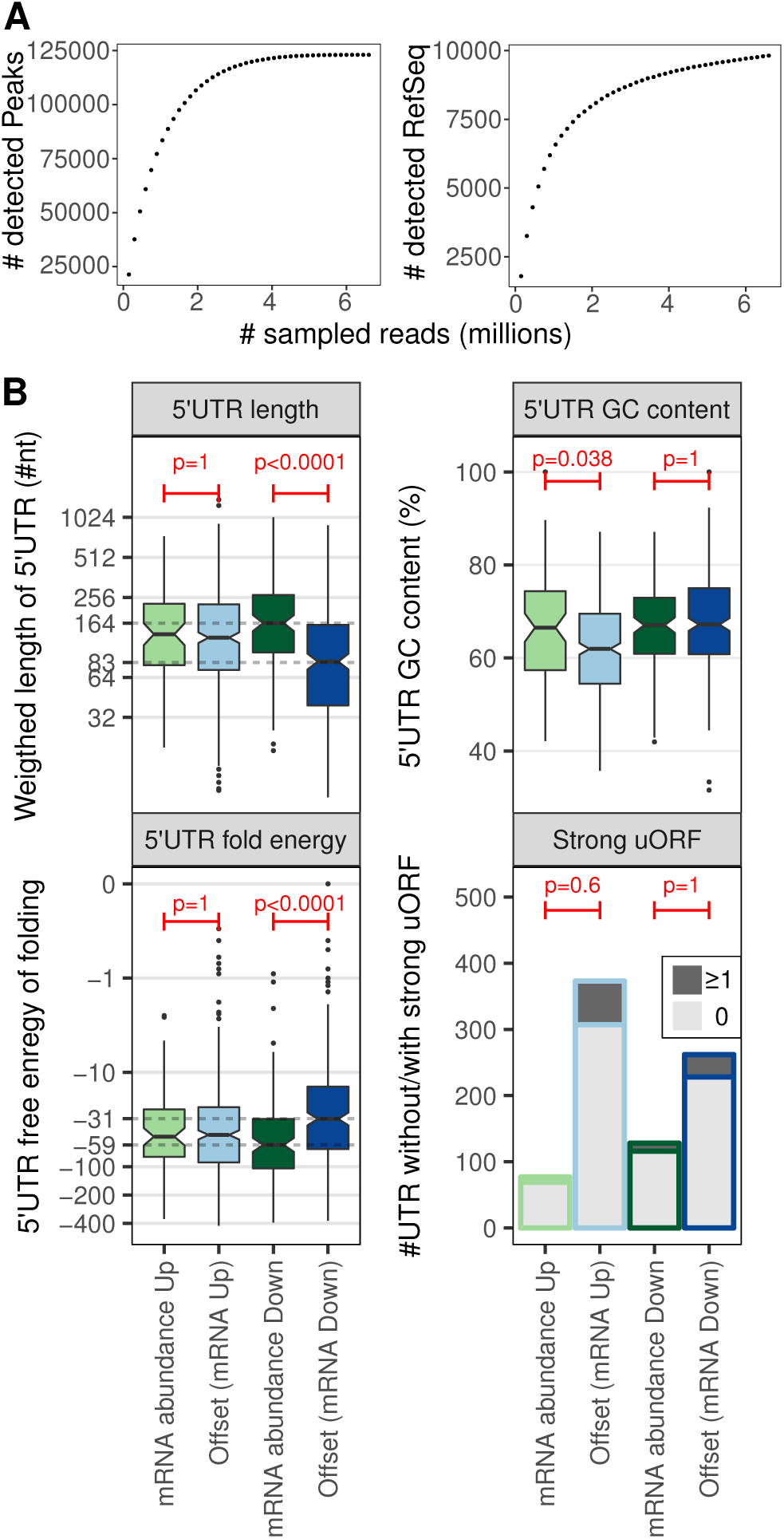
Genes repressed upon ERα depletion but translationally offset have short and less stable 5’ UTRs. **(A)** nanoCAGE sequencing was applied to determine transcription start sites in shERα BM67 cells. The number of detected transcription start sites (peaks) and RefSeq transcripts when sampling increasing number of RNAseq reads are indicated to evaluate the complexity of nanoCAGE RNAseq libraries. **(B)** Boxplots for offset and non-offset mRNAs comparing 5’ UTR weighted lengths (log2 scale), GC content (%), free energy (log10 scale, kcal/mole). Number of transcripts harboring at least one (dark gray) or no (light gray) uORF in a strong Kozak context.

### Transcripts whose levels are downregulated but translationally offset upon ERα depletion are largely devoid of miRNAs target sites

ERα modulates expression of multiple miRNAs (Castellano *et al*, 2009; Maillot *et al*, 2009; Klinge, 2012; Bailey *et al*, 2015) which led us to investigate the role of miRNAs in translational offsetting. To this end, we performed small RNAseq in shERα and control BM67 cells (**Supplementary Fig. S4**). ERα depletion led to alterations in levels of a subset of miRNAs (**Fig. 4A**). Among these, five miRNAs (miR-181a-5p, miR-21a-5p, miR-23b-3p, miR-32-5p, miR27b-3p) were selected and their expression was validated using qPCR (**Fig. 4B-C**). As RNAseq involves relative quantification of miRNAs, we also considered a global change in miRNA expression depending on ERα. Bioanalyzer based quantification of small RNAs to assess global changes in miRNA expression, however, indicated no difference in total miRNA expression between shERα and control cells (**Fig. 4D**). Next, we assessed whether targets of miRNAs with ERα-dependent expression were selectively offset. Transcripts upregulated but translationally offset in control vs shERα BM67 cells showed no enrichment of target sites for downregulated miRNAs (**Fig. 4E-G**). In contrast, downregulated mRNAs that were translationally offset were largely devoid of the target sites for miRNAs which were upregulated in ERα depleted cells (**Fig. 4H-I**). Importantly, such a strong underrepresentation of miRNA target sites among transcripts whose abundance was reduced but translationally offset upon ERα depletion was also observed when selecting random sets of miRNAs (**Fig. 4J**). This suggests that there is no link between ERα-regulated miRNAs and translational offsetting, but rather that there is a general lack of miRNA target sites within this subset of transcripts. In summary, mRNAs whose levels are reduced but offset upon ERα depletion have short and less structured 5’UTRs and harbor less miRNA target sites in their 3’ UTRs as compared to non-offset transcripts.

**Figure 4.**
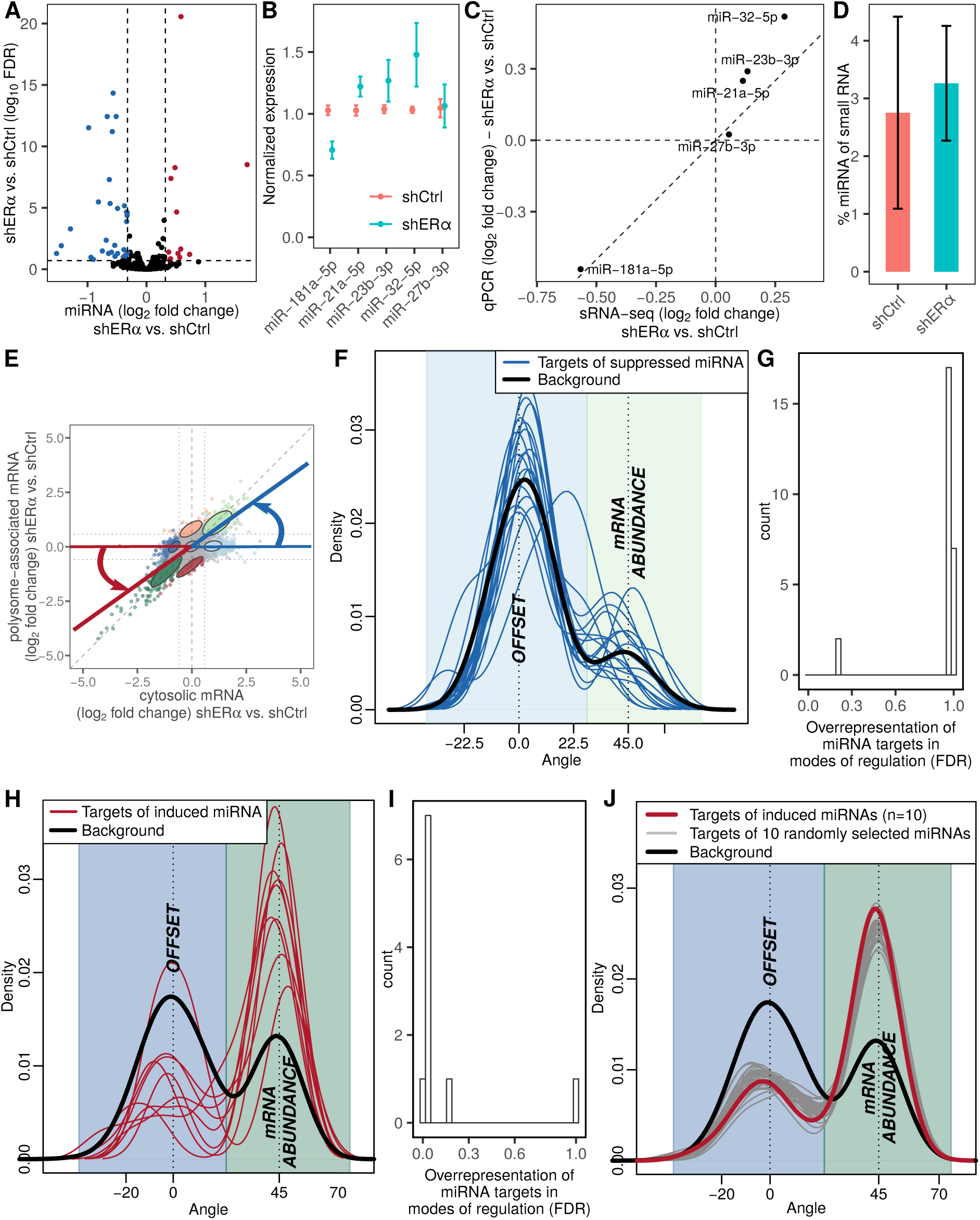
Upon ERα depletion, target sites for miRNAs are under-represented among suppressed but offset mRNAs. **(A)** Volcano plot of ERα dependent changes in miRNA expression. Down- and up-regulated miRNAs are colored in blue and red, respectively. **(B-C)** Validation of miRNA expression using qPCR. Normalized expression (mean ± standard deviation) (B) and expression fold change (shERα vs. shCtrl) obtained from small RNAseq vs. qPCR (C). **(D)** Mean (± standard deviation) of percentage miRNA among small RNAs in shERα and shCtrl BM67 cells. **(E)** Each modulated mRNA was assigned an angle representing mode of regulation as visualized. **(F)** Density of obtained angles for the background (black) and targets of each suppressed miRNA (blue lines). **(G)** Histogram of FDR-adjusted p-values from Fisher’s exact tests assessing over-representation of induced and offset mRNAs vs. induced non-offset transcripts among targets for each suppressed miRNA as compared to non-targets. **(H-I)** same as (F-G) but for suppressed mRNAs and induced miRNAs. (**J**) Density of angles for the background (black), targets of all induced miRNA (red) and targets of randomly selected groups of 10 miRNAs (gray lines).

### Transcripts whose level are induced by ERα depletion, but translationally offset are decoded by a distinct set of tRNAs

Because no distinct 5’ or 3’UTR features were observed among transcripts induced but translationally offset in ERαdepleted vs. control cells, we next investigated their codon usage. Transcripts expressed during proliferation *vs.* differentiation exhibit distinct codon usage and thus appear to require different subsets of tRNAs for their translation (Gingold *et al*, 2014). We therefore considered that a mismatch between tRNA demand (codon usage from expressed mRNAs) and tRNA expression could lead to translational offsetting. Indeed, there was a strong association between the mode of regulation and codon composition (p<0.001, Pearson’s Chi-squared test) as mRNAs whose upregulation was translationally offset following ERα depletion showed a striking enrichment for a distinct subset of codons (**Fig. 5A-B**). This was confirmed by analyses of codon bias using a set of highly expressed genes as reference (top panels) and measures of adaptation to the tRNA pool by the tRNA adaptation index (tAI) whereby relative tRNA levels are assumed to be mirrored by their genomic copy numbers (**Supplementary Fig. S5A**). Moreover, a reduced tAI was observed at all sextiles along the coding sequences of upregulated mRNAs which were translationally offset (**Supplementary Fig. S5B**). We next assessed how codon usage in herein identified modes of regulation of gene expression compared to codon usage in transcripts encoding proteins with distinct cellular function (which was used to detect codon bias between proliferation and differentiation associated-transcripts previously (Gingold *et al*, 2014)). To this end, we first visualized differences in codon usage between mRNAs grouped based on cellular functions (i.e. within GO terms) using correspondence analysis. Codon usage of different modes of regulation of gene expression were then projected in the same dimensions (**Fig. 5C**, **Supplementary Table S3**). Proliferation and differentiation related functions showed extreme positive and negative values, respectively, in the first dimension (**Supplementary Table S3**). Strikingly, mRNAs which were induced at the total transcript level but translationally offset following ERα depletion showed a more positive value in dimension one than any GO term (**Fig. 5C**). Notably, in the gene-set enrichment analysis (**Fig. 2**), while the most enriched GO terms found among proteins encoded by mRNAs whose upregulation was offset at the level of translation were related to cellular proliferation, this enrichment did not pass the thresholds for statistical significance (**Supplementary Table S2**). To further characterize the nature of the differences in codon usage we selected the first quintile of codons (12 codons; **Fig. 5B; Supplementary Table S4**) with highest positive residuals from independence between codon composition and mode of regulation. Out of these, 9 and 3 codons were over-represented among mRNAs whose upregulation and downregulation, were respectively translationally offset, following ERα depletion (**Figs. 5B, 5D)**. Notably, these 3 codons showed a stronger depletion among mRNAs that were upregulated but translationally offset as compared to their enrichment among mRNAs which were downregulated but offset. Yet, the codon adaptation indexes of mRNAs whose downregulation was offset at the level of translation were all consistently higher compared to those of non-offset mRNAs (**Fig. 5B; Supplementary Fig. S5AC; Supplementary Table S4**). These analyses were based on codon frequency (which is affected by amino acid frequency) but comparable results were obtained following normalization to amino acid counts (**Fig. 5E**, **Supplementary Fig. S5D-F**). Notably, codons enriched among induced and offset (9 codons) or repressed and offset (3 codons) mRNAs show strong co-variation across expressed transcripts (**Supplementary Fig. S5G**). In summary, mRNAs whose total levels are induced but offset at the level of translation show distinct codon usage, which suggests that their translational offset could stem from a mismatch between tRNA expression and demand.

**Figure 5.**
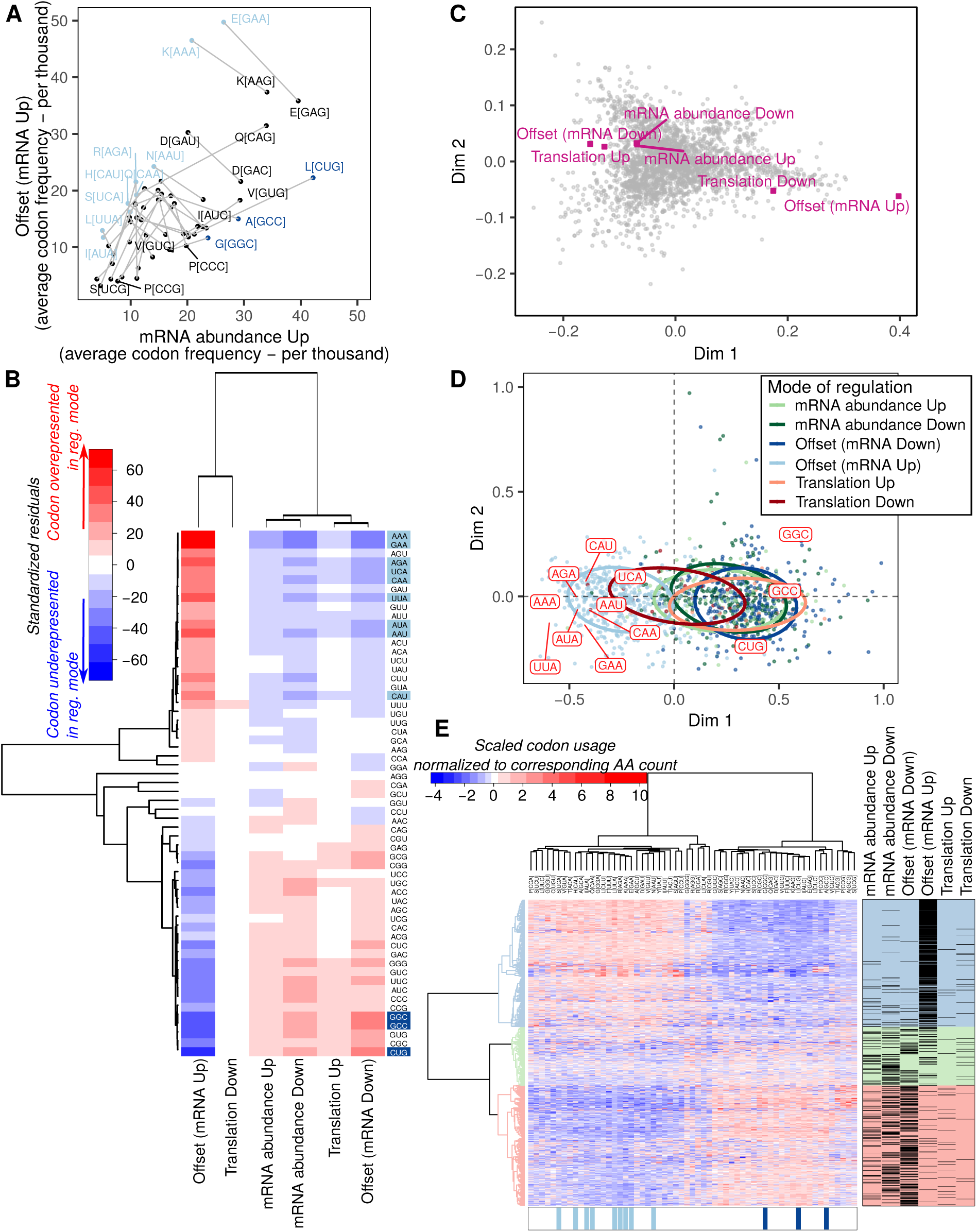
Transcripts induced but translationally offset upon ERα depletion are characterized by distinct codon usage. **(A)** For each codon, the average frequency (per thousand) is compared between transcripts induced but offset vs non-offset (i.e. abundance mode of regulation) upon ERα depletion. Codons for the same amino acid are connected by a gray line. **(B)** Heatmap of standardized residuals from a chi-squared contingency table test. Shown in red and blue are cells with counts significantly higher and lower, respectively, than expected counts under the null hypothesis (i.e. independence between mode for regulation of gene expression and codon composition). **(C)** A correspondence analysis of codon composition for all mouse GO terms. Modes of regulation are projected on the same dimensions. **(D)** A correspondence analysis of codon composition for all regulated mRNAs (from Fig. 1E). mRNAs are projected on the 2 first dimensions of the correspondence analysis and colored according to their mode of regulation. Codons selected as over-represented among translationally offset mRNAs are projected in the same dimensions. **(E)** Unsupervised clustering of gene level codon usage normalized by amino acid counts. All regulated mRNAs (from Fig. 1E) are shown in rows and all codons in columns. Codons identified as over-represented among mRNAs whose levels were induced but offset or suppressed but offset are indicated in light and dark blue, respectively.

### Transcripts whose upregulation is translationally offset are enriched in codons depending on U34-modified tRNAs for their efficient decoding

To explore whether ERα-dependent alterations in tRNA expression may underpin translational offsetting, we employed RNAseq of small RNAs. Notably, tRNA modifications result in short RNA sequencing reads, which do not consistently allow locus-specific expression data (Cozen *et al*, 2015). Nevertheless, we obtained expression data on tRNAs irrespective of loci which allowed quantification of tRNA expression (**Supplementary Fig. S6**). When comparing shERα to control BM67 cells, however, no significant change in tRNA expression was observed (**Fig. 6A**). We next grouped tRNAs corresponding to codons identified as overrepresented in mRNAs whose alterations were translationally offset. No differences in expression of any of the tRNA groups were however observed between shERα vs. control BM67 cells (**Fig. 6B**). In addition to alteration of their expression, tRNA function is regulated by post-transcriptional modifications (El Yacoubi *et al*, 2012).

**Figure 6.**
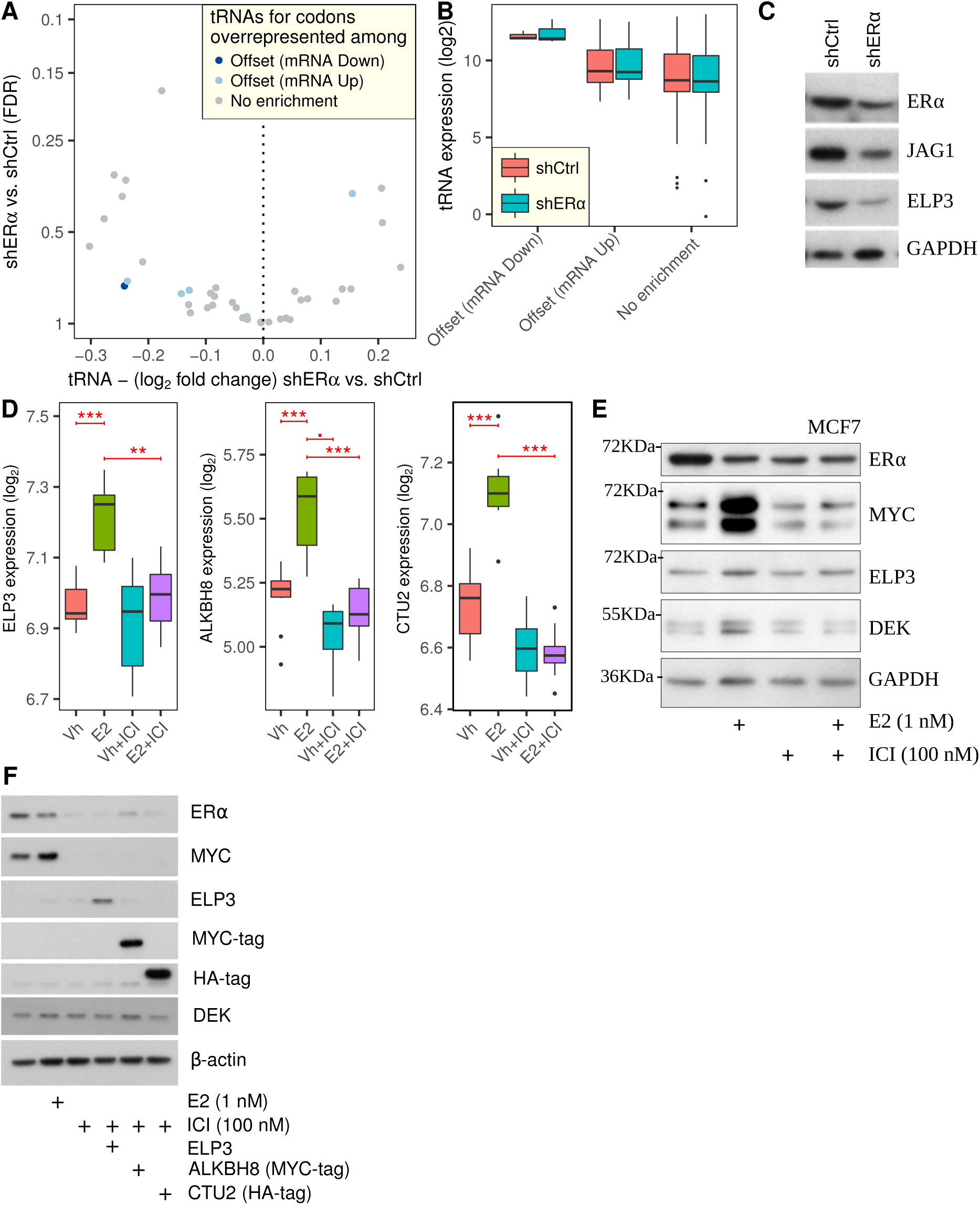
Translational offsetting of transcripts induced following ERα depletion is mediated by U34 tRNA-modification enzymes. **(A)** Volcano plot of changes in tRNA level (shERα vs. shCtrl). Each tRNA is colored according to the mode for regulation of gene expression it is enriched in. **(B)** tRNAs were grouped according to the mode for regulation of gene expression they are enriched in and their expression is compared using boxplots. **(C)** Immunoblotting of BM67 cell extract for ELP3 and JAG1 after transient ERα knockdown in BM67 cells. **(D)** Boxplots of gene expression of ELP3, ALKBH8 and CTU2 upon treatment with 17β-E2 and/or ICI in MCF7 cells (extracted from (Wardell et al, 2012)). *** p<0.001; ** p<0.01; • p<0.1. **(E)** Immunoblotting of MCF7 cell extracts after treatment with 1 nM E2 and/or 100 nM ICI after 24 hours. **(F)** Immunoblotting of MCF7 cell extract after overexpression of ELP3, ALKBH8 and CTU2 combined with treatment of 1 nM E2 and/or 100 nM ICI. Vh, Vehicle; ICI, ICI-182780.

Modifications present at the anticodon loop affect translational rates, sometimes in a codon-specific manner, by modulating the stability of codon-anticodon pairing, and thus limiting decoding to specific nucleotides in wobble position (Rezgui *et al*, 2013; Deng *et al*, 2015). Among the 9 codons which were over-represented in mRNAs whose upregulation was translationally offset, 7 had an adenosine in the 3’ position. These codons are decoded by tRNAs harboring modified uridine at position 34 (U34; **Supplementary Table S4**). Moreover, *Dek* mRNA, which encodes a tumor-promoting protein and whose upregulation is offset at the level of translation upon ERα depletion (**Fig. 1G**), is dependent on 5-methoxycarbonyl-methyl-2-thiouridine (mcm^5^s^2^-U) modification at U34 of corresponding tRNAs (Delaunay *et al*, 2016). In yeast, this modification is present on tRNA^UUC^_Glu_, tRNA^UUU^_Lys_ and tRNA^UUG^_Gln_, wherein it facilitates decoding of GAA, AAA and CAA codons (Johansson *et al*, 2008). Strikingly, these codons are among those that were over-represented in mRNAs whose induction is translationally offset in ERα depleted cells (**Fig. 5B**). Thus, we speculated that although tRNA expression did not change following ERα depletion, alterations in mcm^5^s^2^-U modifications may underlie translational offsetting of mRNAs whose levels are induced.

### ERα regulates expression of tRNA U34 modification enzymes leading to selective translational offsetting

The mcm^5^s^2^-U modification is catalyzed by a cascade of enzymes including ELP3, ALKBH8 and CTU1/2 (Rapino *et al*, 2017). Upon ERα depletion, *Elp3* (p=0.03) and *Alkbh8* (p=0.16) showed reductions in their amounts of polysome-associated mRNA and, consistently, ELP3 protein level was decreases in shERα BM67 as compared to control cells (**Fig. 6C**; no reagents were available to measure mouse ALKBH8). There are two approaches to studying estrogen signaling, depleting the receptor or modulating its activity with ligands. Using BM67 cells with stable knockdown of ERα excluded any confounding effects of ERβ and established the translational offsetting as a sustained response. Yet, this model does not distinguish between direct effects of ERα –which are usually observed after 2 to 24 hours of estradiol treatment–and indirect regulation induced by its transcriptional targets (Katchy *et al*, 2012). We therefore sought to independently confirm these findings using MCF7 human breast cancer cells, one of the most common models of ligand-induced ERα activity (Hewitt & Korach, 2018). To this end, we used a data set where human breast cancer MCF7 cells were starved followed by treatment with estradiol (E2) or vehicle for 24 hours in the presence or absence of selective estrogen receptor modulators (SERMs) (Wardell *et al*, 2012). Indeed, this revealed that *ELP3, ALKBH8* and *CTU2* mRNA levels are induced in an ER-dependent fashion (**Fig. 6D**; *CTU1* was not quantified). As previously reported, stimulation of MCF7 cells with E2 leads to reduced ERα protein expression due to ligand-receptor induced conformational change and poly-ubiquitination following stimulation and receptor activation (Wijayaratne & McDonnell, 2001) (**Fig. 6E**). Transcriptional activity of ERα was confirmed by upregulation of MYC (Wang *et al*, 2011) which was paralleled by increase in ELP3 and DEK protein levels (**Fig. 6E**). This response was abrogated by ER antagonist ICI-182780 (**Fig. 6E**). This suggests that translational offsetting of mRNAs whose levels are induced by ERα depletion may be mediated by ERα-dependent alterations in mcm^5^s^2^-U modification. Notably, as observed in a recent study (Rapino *et al*, 2018), the regulation of these enzymes did not correlate with global changes in mcm^5^s^2^-U-modified tRNAs as quantified by the [(N-acryloylamino)phenyl]mercuric chloride (APM) method (Igloi, 1988) (**Supplementary Fig. S7**). Nonetheless, we sought to rescue E2-stimulated DEK expression by overexpressing the mcm^5^s^2^-U34 modification enzymes in MCF7 cells which were treated with E2 and/or ICI-182780 (**Fig. 6F**). Overexpression of AKLBH8, but not ELP3 or CTU2, rescued DEK protein levels. (**Fig. 6F, Supplementary Fig. S8**). These data suggest that enzymes catalyzing mcm^5^s^2^-U tRNA modifications require ERα and E2 for their expression. Thus, downregulation of U34-modifying enzymes following ERα-depletion leads to translational offsetting of upregulated transcriptional targets.

## Discussion

Improved experimental and analytical methods for transcriptome-wide analysis of translation have been essential for identifying hitherto unprecedented mechanisms of translation regulation (Truitt & Ruggero, 2016; Ingolia *et al*, 2018; Yordanova *et al*, 2018). Using such methods, upon depletion of ERα in prostate cancer, we observed that translational offsetting appears to be a pervasive mechanism which maintains proteome composition. Transcription start site profiling and sequencing of small RNAs revealed that translational offsetting of mRNAs whose levels are decreased is linked to distinct 5’ and 3’ UTR features. The length of the 5’UTR can have strong effects on translation where short (e.g. <30 bases) and very long (e.g. >150 bases) 5’UTRs are associated with reduced translational efficiencies (Pelletier & Sonenberg, 1987; Koromilas *et al*, 1992; Arribere & Gilbert, 2013; Sinvani *et al*, 2015; Gandin *et al*, 2016b). In contrast, the median 5’UTR length (85 nt) of mRNAs whose downregulation is offset at the level of translation corresponds to what has been described as the “optimal” length for translation in mammalian cells (Kozak, 1987). Moreover, target sites for miRNAs, which mediate translational suppression (Filipowicz *et al*, 2008), are largely absent in mRNAs whose suppression is translationally offset. Finally, tRNAs required for decoding mRNAs whose downregulation is translationally offset appear to be expressed at higher levels as compared to other identified tRNA groups (Fig. 6B) and such transcripts are also better adapted to the tRNA pool as compared to non-offset mRNAs (Supplementary Fig. S5A-B). Therefore, this subset of mRNAs exhibits multiple features which would be expected to facilitate translational offsetting upon reduced mRNA levels.

We also observed widespread translational offsetting for mRNAs whose levels were induced in ERα depleted vs. proficient cells. In this case, translational offsetting may be attributed to codon usage of these transcripts. Indeed, the frequency of codons which are decoded more efficiently by the U34-modified tRNAs was substantially higher in transcripts whose induction after ERα depletion was translationally offset as compared to non-offset mRNAs. Consistently, ERα depletion reduced expression of U34 modification enzymes, which appear to play a major role in tumorigenesis (Ladang *et al*, 2015; Delaunay *et al*, 2016; Rapino *et al*, 2018). In this context, genes such as DEK (Delaunay *et al*, 2016) and HIF1α (Rapino *et al*, 2018) were characterized as key downstream effectors, which mediate the pro-neoplastic effects of U34 modifications. Consistently, we demonstrate that DEK expression is induced by ERα and E2. Collectively, these findings suggest that ERα-dependent modulation of U34-modification enzymes expression results in translational offset of transcripts whose levels are induced upon ERα depletion.

In addition to translation initiation, it has recently been revealed that translation elongation is also dysregulated in cancer, which in part appears to be determined by codon composition of mRNAs (Leprivier *et al*, 2013; Faller *et al*, 2015). Herein, we have identified regulation of U34modification enzymes as a process by which ERα modulates translation of a subset of mRNAs with distinct codon usage in prostate cancer cells. This regulation is mediated at the level of mRNA translation by offset of transcriptional targets requiring U34-modified tRNAs. At the same time, a second set of transcripts which harbor optimal 5’UTRs and codon composition but lack target sites for miRNAs are translationally offset when mRNA levels decrease. We speculate that this plays a role in mediating biological effects of ERα in neoplastic tissues. Indeed, using a polysome-profiling data set comparing tamoxifen-sensitive vs. resistant cells (Geter *et al*, 2017), we observed translational activation of mRNAs with higher requirements for U34modified tRNAs and increased expression of ALKBH8 in tamoxifen-resistant cells (**Supplementary Fig. S9**). Thus, modulation of translation via ERα-dependent changes in U34-modifications may be associated with drug resistance and our findings therefore may have important implications in understanding alterations in gene expression programs following treatment with ERα antagonists.

In conclusion, this study establishes translational offsetting as a distinct subtype of a wide-spread buffering mechanism which allows adaptation to acute (E2 treatment) and chronic (ERα depletion) alterations in ERα-dependent transcriptomes. Moreover, these findings unravel a previously unappreciated cooperation between transcriptional and translational programs which suggest a hitherto unappreciated plasticity of gene expression machinery in shaping adaptive proteomes.

## Methods

### Cell Culture, antibodies, western blot

MCF7 cells were purchased from America Type Tissue Culture Collection and used at low passage for less than 2 months before thawing a new vial. The PTEN-deficient prostate cancer cell line (BM67) derived from the PBCre;Pten^Flox/Flox^ mouse model of prostate cancer (Wang *et al*, 2003) has been described previously (Takizawa *et al*, 2015) and used at low passage. Stable knockdown of ERα was achieved using the pGIPZ vector (Open Biosystems) containing a non-silencing control (shCtrl) or a mouse ERα shRNA (V2LMM_30677). Cells were maintained on 4 µg/mL of puromycin (Sigma). All cells were grown in RPMI-1640 (Gibco) and supplemented to a final concentration of 5% Fetal Bovine Serum (FBS, Sigma) and 100 IU/mL penicillin and 10 µg/mL streptomycin (P/S, Invitrogen) and kept in a humidified incubator at 37°C supplemented with 5% CO_2_. All cell lines were routinely tested for mycoplasma (in house service, Peter MacCallum Cancer Centre). Western blot information is provided in **Supplementary Methods**.

### RNA-extraction

Polysome profiling was performed on four replicates of each condition as previously described (Gandin *et al*, 2014). Briefly, cytosolic lysates were loaded on a 5-50% sucrose gradient allowing for isolation of mRNAs associated with more than 3 ribosomes (hereafter referred to as polysome-associated mRNA) after ultracentrifugation. Total cytosolic RNA was isolated in parallel (Gandin *et al*, 2014).

### DNA-microarray assays and data processing

Cytosolic and polysome-associated RNA were quantified using the Affymetrix Mouse Gene 1.1 ST array as described previously (Gandin *et al*, 2016a) by the Bioinformatics and Expression analysis core facility at Karolinska Institutet. Poor quality arrays were obtained for one replicate of each condition leading to exclusion of the whole replicate (cytosolic and polysome-associated samples from both conditions) from all analyses. Gene expression was normalized using Robust Multiarray Averaging and annotated with a custom probeset definition (mogene11st_Mm_ENTREZG) (Dai *et al*, 2005; Sandberg & Larsson, 2007).

### Analysis of polysome-profiling data

Changes in translational efficiency leading to altered protein levels were quantified using analysis of partial variance (Larsson *et al*, 2010, 2011) as implemented in the anota2seq R/Bioconductor package version 1.2.0 (Oertlin *et al*, 2018). Differential expression of cytosolic and polysome-associated RNA was also assessed using the anota2seq package. For such analyses, Benjamini-Hochberg correction was used to account for multiple testing and a random variance model was used to increase statistical power (Wright & Simon, 2003; Larsson *et al*, 2010). Translational offsetting was defined by mRNAs showing changes in cytosolic RNA which are not reflected in polysome loading as implemented in anota2seq. Details of the analysis are provided in **Supplementary Methods**.

### Nanostring gene expression quantification and analysis

145 genes were selected for quantification by Nanostring (Geiss *et al*, 2008). Within each mode of regulation (as defined in Fig. 1E), genes among the top smallest p-values were randomly selected. Additional genes within each mode of regulation (not belonging to the most significant sets) as well as 11 negative controls (non-regulated genes) were included as well. Details about methodology and analysis are given in **Supplementary Methods**.

### GO enrichment analysis

A Generally Applicable Gene-set Enrichment for Pathway Analysis (Luo *et al*, 2009) was performed to identify enrichment of key cellular functions represented by GO terms (Gene Ontology Consortium, 2015). Additional information is provided in **Supplementary Methods**.

### nanoCAGE library preparation, sequencing and analysis

nanoCAGE libraries of cytosolic mRNA from shERα BM67 cells were prepared as described previously (Gandin *et al*, 2016b) with several modifications detailed in **Supplementary Methods** where description of pre-processing and analysis is also provided.

### RNAseq of small RNAs

RNA was extracted in triplicates from ERα shRNA and control BM67 cells using the RNeasy Plus Mini Kit (Qiagen). RNAseq libraries were prepared according to the Illumina TruSeq Small RNA Library Preparation protocol with small RNA enrichment on the Agilent Bravo Liquid Handling Platform and sequenced on HiSeq2500 (HiSeq Control Software 2.2.58/RTA 1.18.64) with a 1×51 setup. Library preparation and sequencing was performed at Science for Life Laboratory National Genomics Infrastructure. Preprocessing and analysis of RNAseq of small RNAs data are described in **Supplementary Methods.**

### Validation of miRNA expression using qPCR

cDNA was synthesized using miSCRIPT II RT kit (Qiagen; 3 replicates) followed by specific miRNA amplification using miSCRIPT SYBR Green PCR kit (Qiagen) using the following miScript specific primers mmu-miR-21a-5p (5’-UAGCUUAUCAGACUGAUGUUGA-3’), mmu-miR-181a5p (5’-AACAUUCAACGCUGUCGGUGAGU-3’), mmu-miR-32-5p (5’-UAUUGCACAUUACUAAGUUGCA-3’), mmu-miR-23b-3p (5’-AUCACAUUGCCAGGGAUUACC-3’) and mmumiR-27b-3p (5’-UUCACAGUGGCUAAGUUCUGC-3’).

### Analysis of codon usage

A detailed explanation is given in **Supplementary Methods**. Briefly, the longest coding sequences of all regulated mRNAs were extracted from the consensus coding sequence database (Pruitt *et al*, 2009) to retrieve their codon composition. The codon usage indexes were computed using the codonW (CodonW) and tAI (dos Reis *et al*, 2003, 2004; dos Reis) packages.

### Quantification and analysis of tRNA levels and modifications

RNAseq of small RNAs data described above was used for tRNA quantification using the ARM-seq bioinformatics pipeline from Cozen et al. (Cozen *et al*, 2015) with a few modifications detailed in **Supplementary Methods**. For quantification of mcm^5^s^2^-U modified tE(UUC), tRNA was purified using the miRvana kit (Roche; 4 replicates). 0.5µg RNA was resolved on an 8%. acrylamide gels containing 0.5x TBE, 7 M urea, and 50 mg/mL [(N-acryloylamino)phenyl]mercuric chloride (APM) (Igloi, 1988). Northernblot analysis was performed essentially as described in Leidel et al. (Leidel *et al*, 2009), using 5’-TTCCCATACCGGGAGTCGAACCCG-3’ as probe to detect tE(UUC).

### Analysis of public dataset for E2 dependent expression of ELP3, ALKBH8 and CTU2

Expression of enzymes catalyzing the mcm^5^s^2^-U34 tRNA modification upon treatment with E2 and/or ICI-182780 was analyzed using a publicly available data (Wardell *et al*, 2012) as detailed in **Supplementary Methods**.

### Selective Estrogen Receptor Modulator (SERM) treatment and target gene rescue

Cells were seeded in standard cell culture conditions to achieve 25-40% confluency. Twenty-four hours later, the culture medium was replaced with fresh standard culture medium (FBS). 24 hours later, SERM were administered at a concentration of 1 nM 17β-Estradiol (E2) or 100 nM of Fulvestrant (ICI-182,780). For target gene rescue, an additional timepoint was added. 24 hours later, samples were incubated with a mixture containing 7.5 µL of Lipofectamine 3000 (Invitrogen) complexed with 2.5 ug of empty vector plasmid or plasmid harboring of ELP3 (pCS6-ELP3, Integrated Sciences #TCMS1004), CTU2 (pcDNA3.1+CTU2-C-HA, GenScript Clone ID OMu03000C) or ALKBH8 (pcDNA3.1+ALKBH8-C-Myc, GenScript Clone ID OMu01949C). Samples were then processed for western blot.

### Statistics

Unless stated otherwise, all statistical tests are two-sided.

### Data availability

The DNA-microarrays, RNAseq of small RNAs and nanoCAGE datasets generated and analyzed during the current study are available in the NCBI Gene Expression Omnibus repository under accession number GSE120917 (https://www.ncbi.nlm.nih.gov/geo/query/acc.cgi?acc=GSE120917).

## Acknowledgements

The authors would like to acknowledge support from Science for Life Laboratory, the National Genomics Infrastructure (NGI) and Bioinformatics and Expression Analysis (BEA, Karolinska Institutet) core facilities and Uppmax for providing assistance in massive parallel sequencing, DNA-microarray processing and computational infrastructure. The research was supported by the Swedish Research Council, the Wallenberg Academy Fellow Program, the Swedish Cancer Society, the Cancer Society in Stockholm and STRATCAN (to OL); the Department of Health and Human Services acting through the Victorian Cancer Agency (MCRF16007) and NHMRC grant (APP1141339) (to LF); Canadian Institutes for Health Research (MOP-363027) and National Institutes of Health grants (R01 CA 20202101-A1) (to ITo). ITo is supported by Junior 2 award from Fonds de Recherche du Québec – Santé. This project was further facilitated by funding from the Swedish foundation for international cooperation in research and higher education (STINT) to ITo, OL and LF.

## Author contributions

MGL, ITa, SAB, GPB, ITo, OL, LF designed the study; RJB, VH, MGL, EK, BS, PB, KS performed experiments; JL, VH, KJS, OL performed analysis and interpreted the data; MGL, ITa, SAB, GPB, ITo, OL, LF supervised the research. All authors edited and approved the final manuscript.

