## Supplementary figures, table legends and methods for "Estrogen receptor alpha controls gene expression via translational offsetting"

Supplemental Fig S1

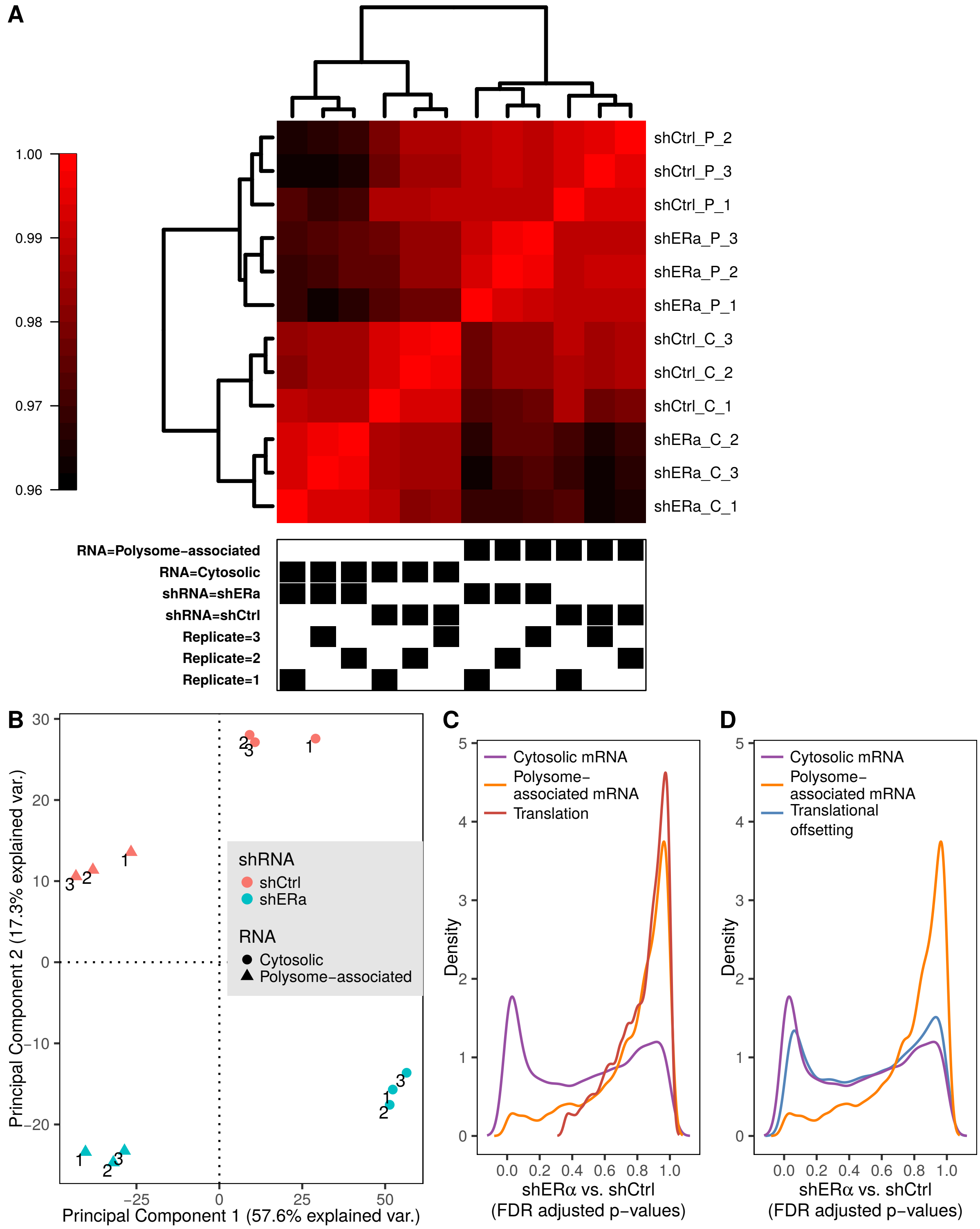


***Figure S1. Quality control for data set assessing transcriptome-wide effects on gene expression programs using polysome-profiling.*** *(**A**) Heatmap of a correlation matrix between all samples from the transcriptome wide study (lower correlations in black and higher correlations in red). Samples were reordered according to the hierarchical clustering. (**B**) Samples from the transcriptome-wide polysome-profiling of ERα depletion (normalized data) were projected on the 2 first components of a centered principal component analysis. (**C**) Density of FDR-adjusted p-values for differential expression between shERα and shCtrl BM67 cells using data from polysome-associated mRNA (orange) or cytosolic mRNA (purple) and for analysis of differences in translational efficiencies leading to altered protein levels (red). (**D**) Same plot as in (C) but including a blue density of FDR-adjusted p-values from analysis of translational offsetting.*

Supplemental Fig S2

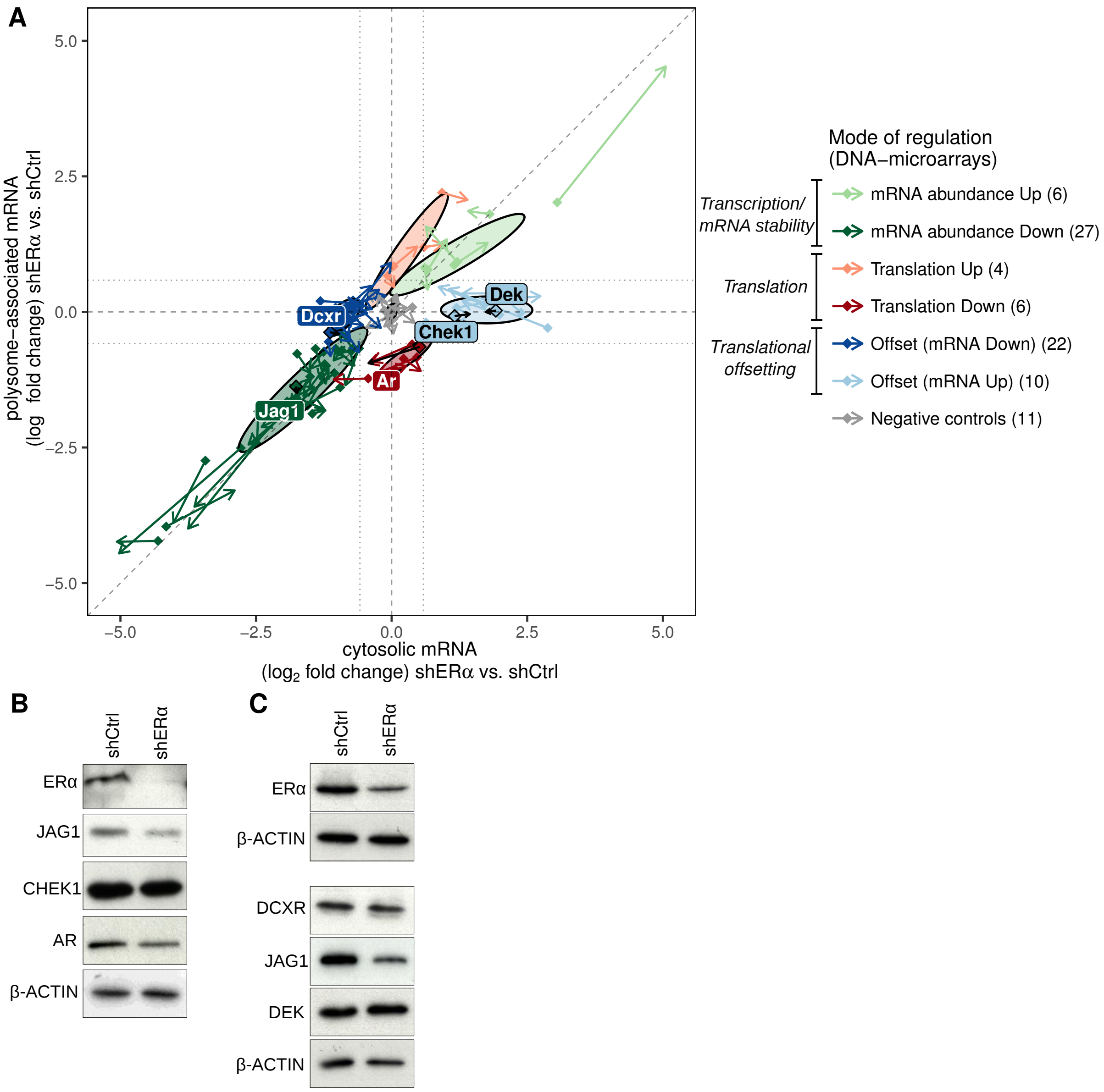


***Figure S2. Validation of translational offsetting by Nanostring and western blotting.*** ***(A)*** *Scatter plot of polysome-associated mRNA vs. cytosolic mRNA* $\log_2$ *fold-change (shERα vs. shCtrl). Confidence ellipses are overlaid (based on fold-changes from* ***Fig. 1E****). For each gene assessed using Nanostring, an arrow is plotted where the start of the arrow indicates fold-changes for the transcriptome wide study (i.e. Fig. 1E) and the tip of the arrow by fold-changes as measured by Nanostring (i.e. Fig. 1F). Genes and arrows are colored according to their mode of regulation.* ***(B-C),*** *Immunoblotting of BM67 cell extract for AR, JAG1, CHEK1, DEK and DCXR after ERα knockdown. Independent experiments from Fig. 1G.*

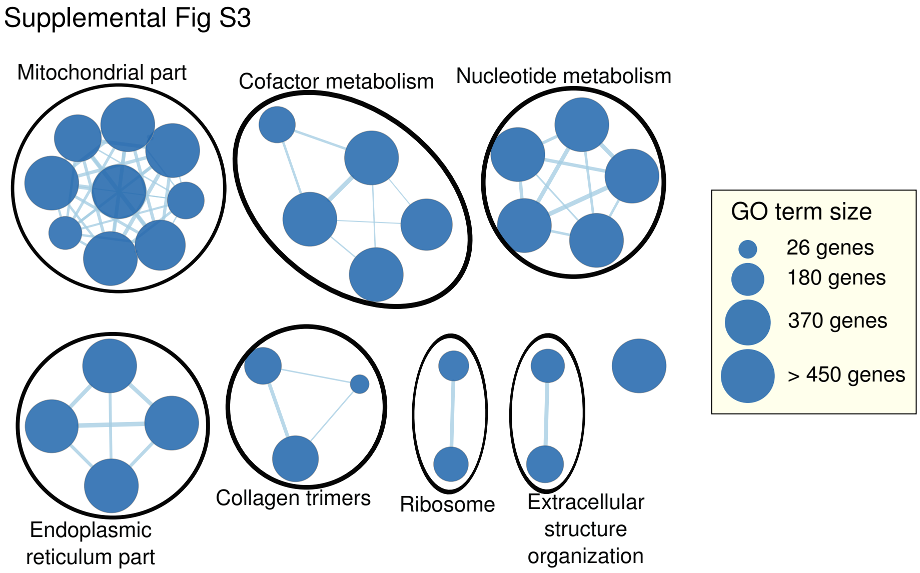


***Figure S3. Cellular functions enriched among proteins encoded by mRNAs whose levels are suppressed.*** *GO terms enriched among proteins encoded by cytosolic mRNAs downregulated (FDR < 0.15) in BM67 cells as compared to control are visualized as a network. Each node is a GO term connected to other GO terms whose annotated genes overlap. The size of each node relates to the number of genes associated to its corresponding term and the width of the edge illustrates the size of the overlap between two connected nodes.*

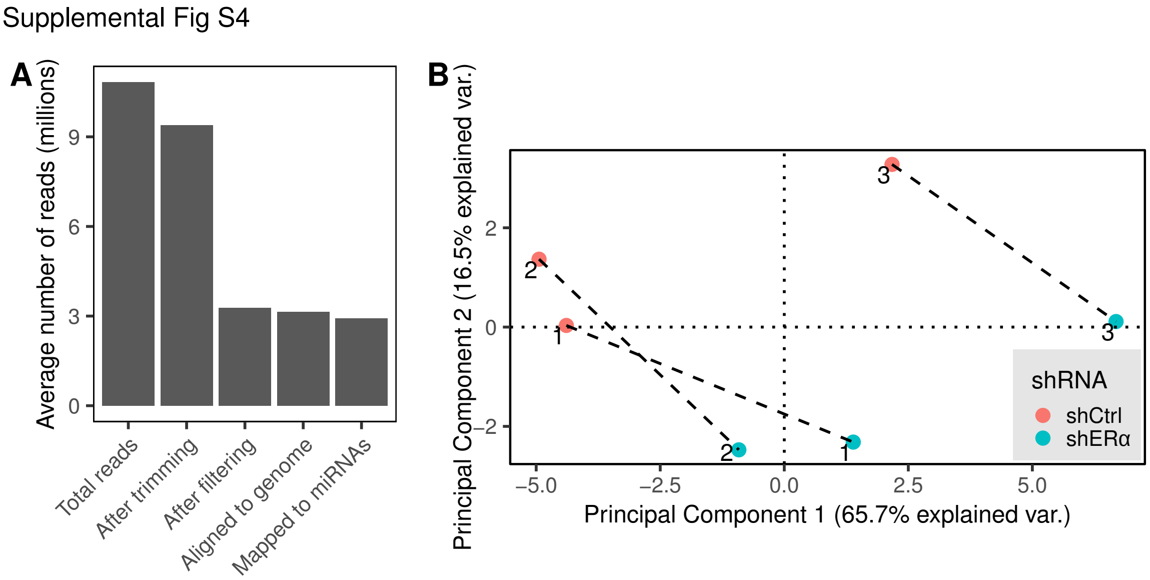


***Figure S4. Quality control for small RNA-seq data on BM67 shERα and control cells.*** ***(A)*** *Average (over the 6 samples) number of reads after each preprocessing step.* ***(B)*** *Triplicate samples (1-3) from the small RNA-seq data were projected on the 2 first components of a centered principal component analysis. miRNAs with less than 30 counts on average have been excluded and the data were normalized using DESeq2 size factors.*

Supplemental Fig S5

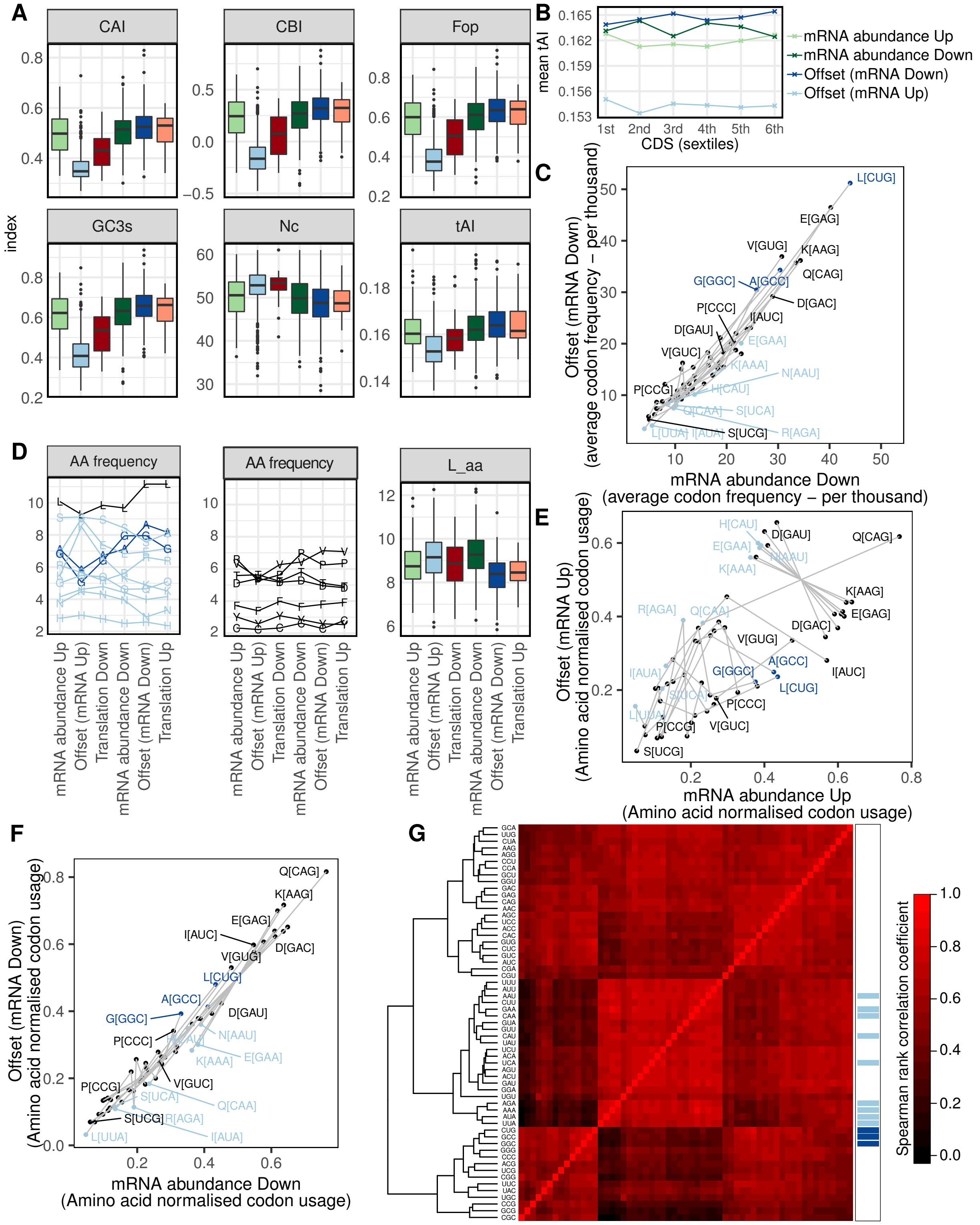


***Figure S5. Analyzes of codon usage.*** ***(A)*** *Boxplots of mRNAs under all modes for regulation of gene expression comparing 6 measurements of codon usage: Codon Adaptation Index (CAI), Codon Bias Index (CBI), Frequency of optimal codons (Fop), GC content at the codon third position (GC3s), Effective number of codons (Nc), tRNA adaptation index (tAI).* ***(B)*** *CDS are divided in sextiles and the mean tRNA adaptation index is plotted for each mode of regulation at these CDS positions.* ***(C)*** *For each codon, the average frequency (per thousand) is compared between transcripts suppressed but offset vs non-offset upon ERα depletion. Codons coding for the same amino acid are connected by a gray line.* ***(D)*** *Left panel shows average frequency (percent) of amino acids encoded by codons previously identified as over-represented among induced or suppressed but offset mRNAs (colored in light or dark blue, respectively; Leucine (L, black) is encoded by codons in both groups). Middle panel shows average frequency (percent) of all other amino acids. Right panel shows boxplots for all mRNAs within each mode for regulation of gene expression comparing number of amino acids ($\log_2$, L_aa, right panel).* ***(E-F)*** *For each codon, the codon usage normalized by amino acid count is compared between transcripts induced (E) or suppressed (F) but offset vs non-offset (i.e. abundance mode of regulation) upon ERα depletion. Codons coding for the same amino acid are connected by a gray line.* ***(G)*** *Unsupervised clustering of a Spearman correlation matrix of codon frequency of regulated mRNAs. Codons identified as over-represented among induced but offset and suppressed but offset mRNAs are indicated in light and dark blue, respectively.*

Supplemental Fig S6

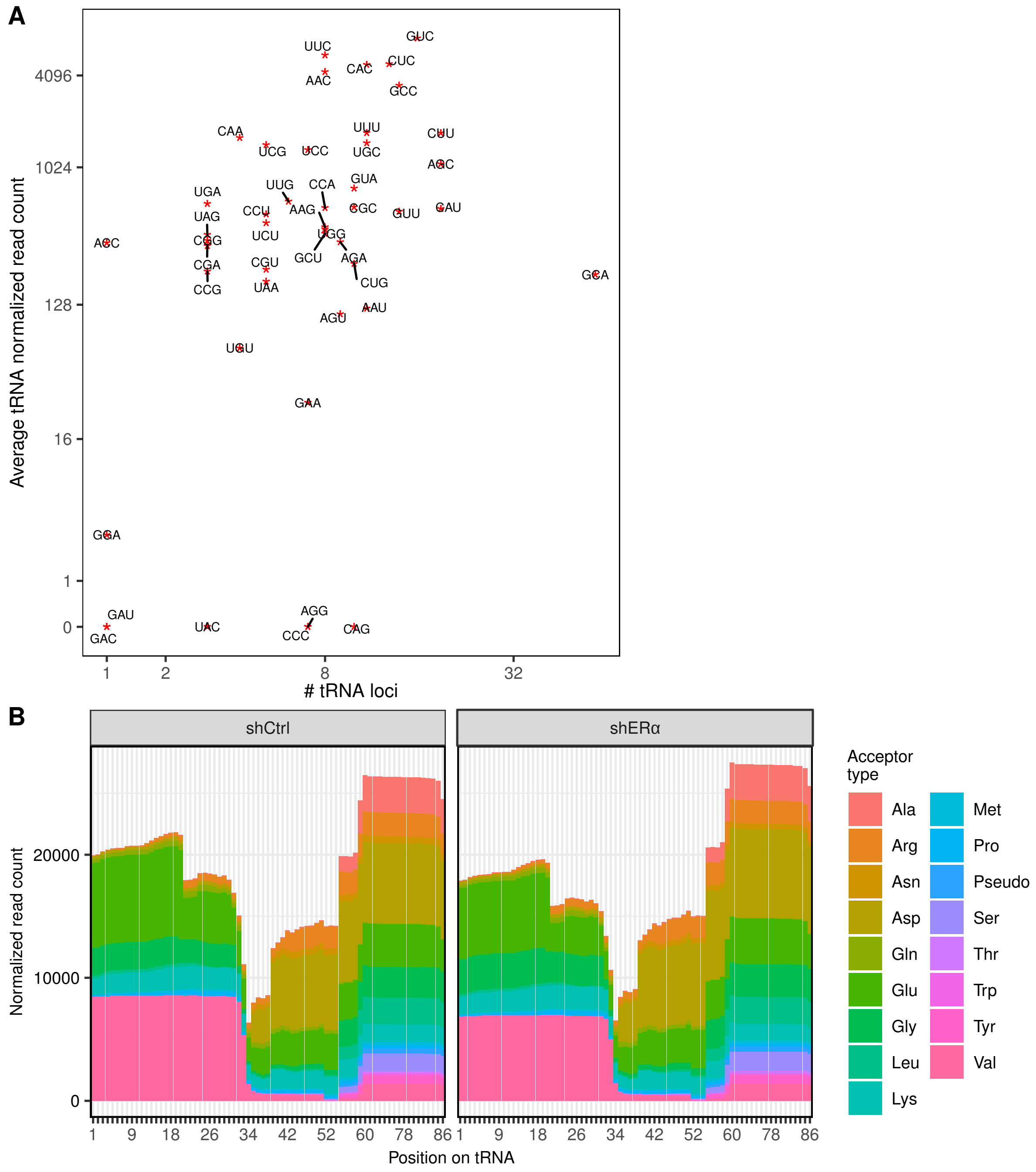


***Figure S6. Quality control for quantification of tRNA levels from small RNA sequencing.*** ***(A)*** *DESeq2 normalized count of all mature tRNAs were summed per anticodon and the average across the 6 samples is plotted against number of tRNA loci (retrieved from UCSC table browser). Quantitative data could be retrieved for most tRNAs (counts > 100),* ***(B)*** *tRNA coverage as measured by the ARM-Seq pipeline. Expected hard-stop modifications can be observed at positions 34 and 58.*

Supplemental Fig S7

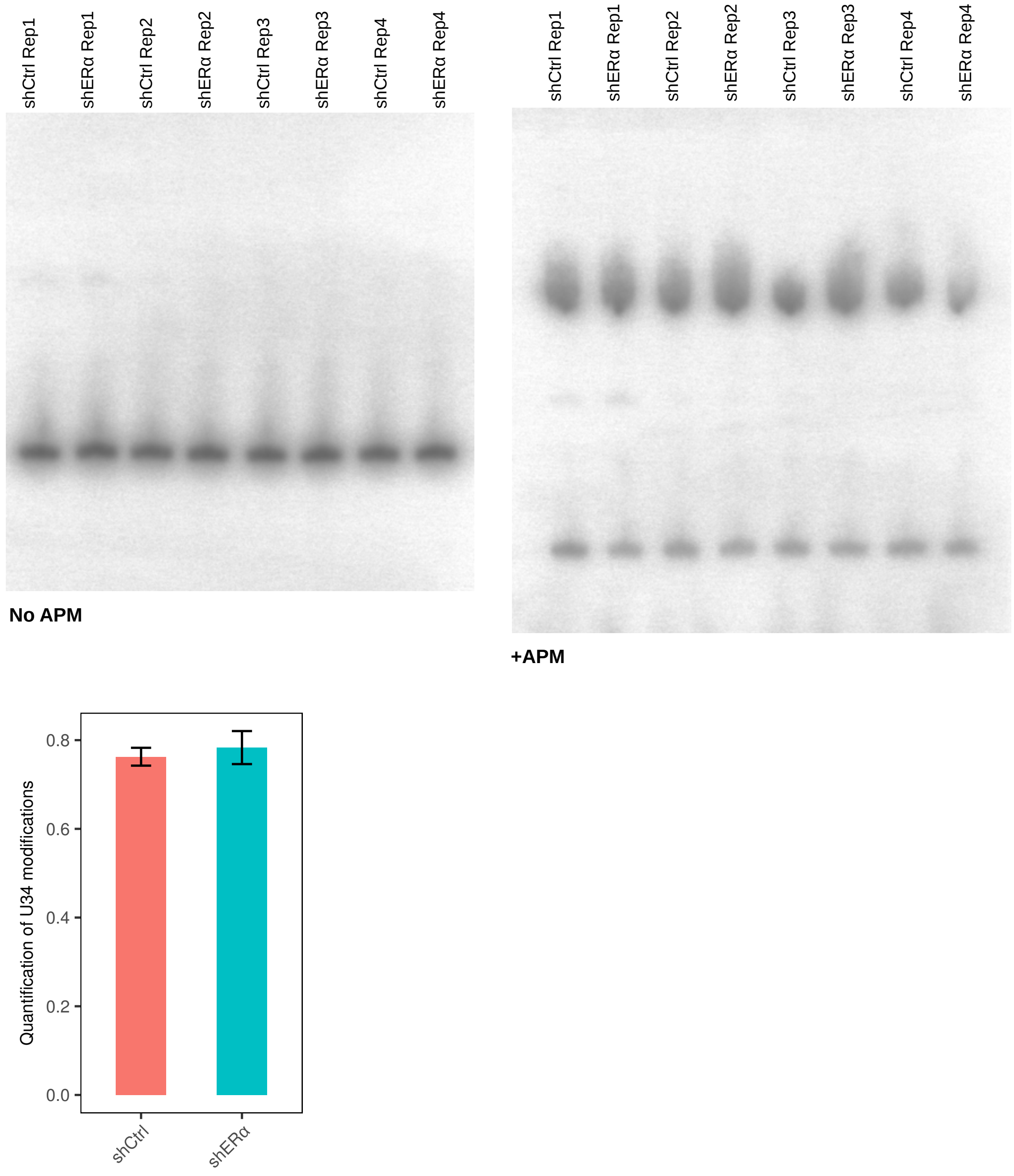


***Figure S7. Quantification of tRNA thiolation.*** *APM-PAGE and subsequent northern blot analysis of tE(UUC) isolated from 4 replicates of BM67 shERα and control cells. Bottom panel shows quantification of the northern blot (mean ± sd of relative amount of modified tRNA as compared to total tRNA; Student t-test* $p = 0.37$*).*

Supplemental Fig S8

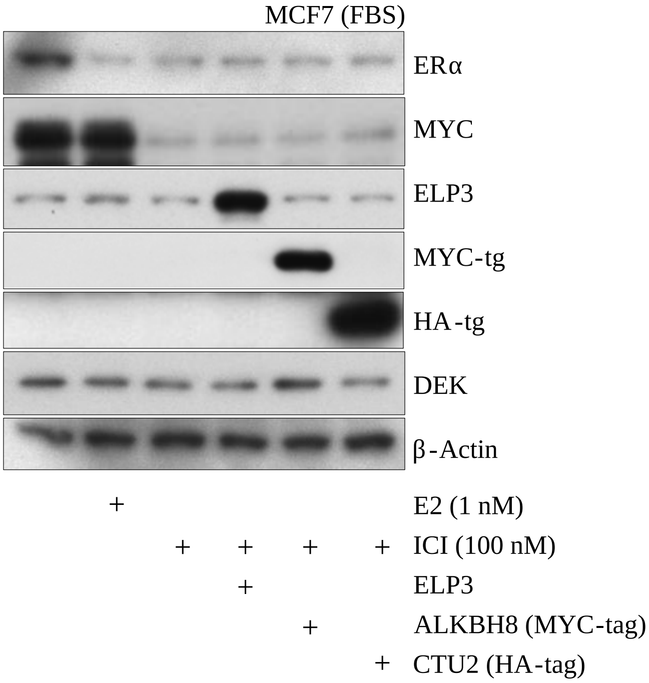


***Figure S8. Modulation of DEK protein expression upon overexpression of U34-modifying enzymes.*** *Immunoblotting of MCF7 cell extract after overexpression of ELP3, ALKBH8 and CTU2 combined with treatment of 1 nM E2 and/or 100 nM ICI. Independent experiment from Fig. 6F. ICI, ICI-182780.*

Supplemental Fig S9

**A**

| | log2 fold change | p-value | FDR-adjusted p-value |
|---|---|---|---|
| ELP3 | -0.06 | 0.74 | 0.87 |
| ALKBH8 | 0.58 | 0.02 | 0.16 |
| CTU1 | 0.08 | 0.69 | 0.84 |
| CTU2 | 0.42 | 0.09 | 0.30 |

**B**

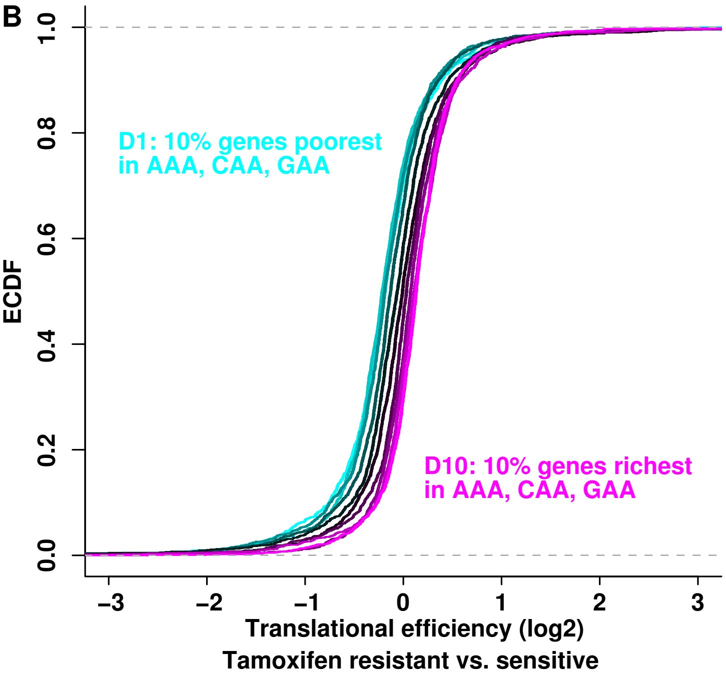


| signature | Wilcox_pval | q25 | q50 | q75 |
|---|---|---|---|---|
| D2 vs. D1 | 3.3e−01 | 0.016 | −0.027 | −0.037 |
| D3 vs. D1 | 6.8e−01 | 0.034 | −0.0042 | −0.013 |
| D4 vs. D1 | 4.6e−05 | 0.11 | 0.057 | 0.031 |
| D5 vs. D1 | 1.8e−16 | 0.16 | 0.12 | 0.095 |
| D6 vs. D1 | 4.9e−33 | 0.21 | 0.18 | 0.17 |
| D7 vs. D1 | 3.7e−52 | 0.29 | 0.22 | 0.2 |
| D8 vs. D1 | 2.9e−72 | 0.32 | 0.26 | 0.24 |
| D9 vs. D1 | 2.3e−87 | 0.35 | 0.3 | 0.28 |
| D10 vs. D1 | 5.5e−98 | 0.39 | 0.32 | 0.26 |

***Figure S9. Translational regulation via the U34-modification pathway may promote resistance to tamoxifen in breast cancer cells.*** **(A)** *Analysis of ELP3, ALKBH8, CTU1/2 expression between Tamoxifen resistant and Tamoxifen sensitive breast cancer cells (data from (Geter* et al*, 2017), see Supplementary Methods). The analysis is performed on polysome-associated mRNA.* **(B)** *Human genes were categorized into deciles of AAA, CAA, GAA codon frequencies in their coding sequences. Cumulative distributions for changes in translational efficiency are plotted for each decile (Tamoxifen resistant vs. tamoxifen sensitive). Statistics are given for the comparison of each decile to the first decile (D1).*

### Supplementary tables

**Supplementary Table S1.** List of transcripts regulated by one of 3 modes of regulation: mRNA abundance (parallel changes in total and polysome-associated mRNA), translation (changes in polysome-associated mRNA without corresponding changes in total mRNA) and translational offsetting (changes in cytosolic mRNA without corresponding changes in polysome-associated mRNA). For each gene and mode of regulation, the $\log_2$ Fold Change (log2FC), unadjusted and FDR-adjusted p-values are provided.

**Supplementary Table S2.** Top 100 GO terms enriched in changes in cytosolic mRNA, polysome-associated mRNA or translational efficiency leading to altered protein levels or offsetting.

**Supplementary Table S3.** GO terms with highest contributions to dimensions 1 and 2 (top 20 GO terms in each direction) of the correspondence analysis represented in Fig. 5C.

**Supplementary Table S4.** Codons identified as enriched in translationally offset mRNAs. The residuals of a test of independence between codon count and mode for regulation of gene expression are given together with the most overrepresented mode of regulation for the first quintile positive residuals. Most of these codons are read by U34 modified tRNAs. The corresponding U34 modification is provided in this table.

### Supplementary methods

#### Analysis of polysome-profiling data

To verify whether the reproducibility of data from polysome-associated mRNA and cytosolic mRNA samples was comparable, we visualized 2 by 2 Pearson correlation coefficients in a clustered heatmap. The clustering was done using default parameters of the annHeatmap2 function (Heatplus 2.26.0 R/Bioconductor package (Ploner, 2015)). We also performed a centered Principal Component Analysis after variance filtering (exclusion of the first quartile).

Anota2seq (Oertlin *et al*, 2018) classifies transcripts into three modes for regulation of gene expression: changes in mRNA abundance (i.e. similar changes in cytosolic and polysome-associated mRNA), changes in translational efficiency leading to altered protein levels, and translational buffering (i.e. offsetting) of changes in cytosolic mRNA levels. The following significance thresholds were used in the anota2seqSelSigGenes function: minEff = $\log_2(1.5)$; maxPAdj = 0.1 (i.e. FDR < 0.1 [for analysis of changes in translational efficiency leading to altered protein levels, no genes passed this threshold. To allow all modes of regulation to be evaluated in downstream analyses, this threshold was relaxed and maxP = 0.05 i.e. p-value < 0.05 was used instead]), selDeltaPT = selDeltaTP = selDeltaP = selDeltaT = $\log_2(1.5)$; maxSlopeTranslation = 1.5; minSlopeTranslation = -1; maxSlopeBuffering = -1.5; minSlopeBuffering = 1. For genes belonging to more than one regulatory mode, a priority order was established such that mRNAs identified as changing their translational efficiency leading to altered protein levels will belong to the translation group and no other group; mRNAs that change their mRNA abundance but are not in the translation group will be in the mRNA abundance group; and buffered (offset) mRNAs are not in the former two groups (implemented in anota2seqRegModes). Genes that were not allocated to any regulatory pattern were called "non regulated". The different regulatory modes are visualized in scatter plots of polysome-associated mRNA $\log_2$ Fold Change (shERα vs. shCtrl) vs. cytosolic mRNA $\log_2$ Fold Change (shERα vs. shCtrl) using confidence ellipses as implemented in the ggplot2 R package (stat_ellipse function) (Wickham, 2009). The smooth scatterplot in Fig. 1C was performed using the stat_density2d function in ggplot2. The plotting area was divided in 16384 squares ($128^2$) of equal size and the density was computed using a two-dimensional kernel density estimation (kde2d; (Venables & Ripley, 2002)). The higher the number of genes there are in each square, the darker the color is on a scale from white to dark blue. Because the estimation is based on smooth distributions, the color of a square also depends on the number of genes in its neighboring squares.

#### Nanostring gene expression quantification and analysis

Nanostring was performed as described previously (William *et al*, 2018) on 3 replicates of cytosolic and polysome-associated mRNA (isolated as described above) from both shERα and control BM67 cells using a custom panel of 145 genes. Position of samples on the lanes of 2 different cartridges were randomized.

The resulting Nanostring counts were $\log_2$ transformed and normalized using the geometric mean of reference genes as normalization factors. Actb, Emc10 are Mcmbp were used as reference genes. Genes for which less than 3 samples were quantified above a background level set at 6 ($\log_2$ scale) were excluded. This resulted in a total of 75 mRNAs belonging to one of the modes of regulation (from Fig. 1E) and 11 pre-selected negative controls. Anota2seq analysis was performed on the normalized Nanostring data with the same parameters and thresholds as for the DNA-microarray analysis with one modification: the cartridge identifier was added as a batch effect in the models.

#### GO enrichment analysis

Gene to GO term associations (retrieved from org.Mm.eg.db_version 3.6.0 (Carlson, 2018)) based only on inference from electronic annotation (IEA) were excluded. GO terms having 5 to 1000 genes were included in the analysis. Genes were ranked based on their signed (by direction of regulation [i.e. up/down]) $\log_{10}$ FDR-adjusted p-values. We used Wilcoxon-Mann-Whitney tests as enrichment tests in both directions separately (i.e. up or down regulation) within the Generally Applicable Gene-set Enrichment Analysis (GAGE) R/Bioconductor package (Luo *et al*, 2009). GO term enrichment results of offset (mRNA down) and cytosolic Down were visualized using the Enrichment Map plug-in in Cytoscape version 3.6.0 (Shannon *et al*, 2003; Merico *et al*, 2010) using the generic analysis and an FDR threshold of 0.15. All other parameters were kept as default. In this network visualization, each node is a GO term connected to other GO terms whose genes overlap. The size of each node reflects the number of genes associated to its corresponding term and the width of the edge illustrates the size of the overlap between two connected nodes. Groups of GO terms were clustered (black circles on Fig. 2 and Supplementary Fig. S3) using the AutoAnnotate plug-in used with default settings (Kucera *et al*, 2016).

#### nanoCAGE library preparation and sequencing

shERα BM67 cells (3 replicates) were grown until 80% confluency before RNA was extracted with TRI-reagent (Sigma-Aldrich, Stockholm, Sweden) and further purified using the RNeasy MinElute Cleanup kit (Qiagen, Sollentuna, Sweden). RNA quantity and quality were determined by Qubit (Life Technologies, Stockholm, Sweden) and Agilent Bioanalyzer (Agilent Technologies, Kista, Sweden), respectively. The integrity number (RIN) of all samples was greater than 8. Reverse transcription was carried out by mixing 1 μl of cytosolic mRNA (75ng) with 1 μl 0.66 M D-Trehalose, 3.3 M of D-sorbitol, 10 μM MS-RanN6 (Sigma-Aldrich; oligo sequences listed below) and an equimolar mixture of two Template Switching Oligonucleotides (TSO; Integrated DNA Technologies, Leuven, Belgium) with different barcodes (ACAGAT and GTATGA). As described previously (Poulain *et al*, 2017), unique molecular identifiers (UMIs) consisting of 8 random nucleotides were included in the sequence of the TSO in order to distinguish multiples copies of a transcript from PCR duplicates. After heat-denaturation for 5 min at 65° C, following components were added to each sample until a total volume of 10 μl: 1x first-strand buffer (Life Technologies, Stockholm, Sweden), 10 mM DTT (Life Technologies, Stockholm, Sweden), 1 M betaine (Sigma-Aldrich, Stockholm, Sweden), 1 mM dNTP mix (TaKaRa, Malmo, Sweden) and 100 units of SuperScript III (Life Technologies, Stockholm, Sweden). Samples were incubated at 25° C for 5 min, 42° C for 60 min, 72° C for 15 min and chilled on ice. Samples were cleaned of primers and enzymes by using AMPure XP magnetic beads (Beckman Coulter, Sweden) according to the manufacturer's instructions and cDNA was eluted in 30 μl nuclease free water.

A diagnostic qPCR to determine the adequate number of temperature cycles was performed prior to the semi-suppressive PCR (ssPCR) needed for cDNA amplification. For this diagnostic qPCR, 8.5 μl of Takara SYBR Premix Ex Taq (1x, TaKaRa, Malmo, Sweden) containing 100 nM of MS-dir 1F and MS-dir 1R each was mixed with 1.5 μl cDNA (in triplicate) and amplified by qPCR (95° C for 5 min, [65° C for 15 s and 68° C for 2 min] x 40 cycles; CFX96 Touch, Bio-Rad, Sweden). For the ssPCR, 20 μl of the purified first-strand cDNA was mixed with 0.5 μl of MS-dir 1F and MS-dir 1R each (100 nM), 4 μl of water and 25 μl of Kapa HiFI HotStart Ready Mix (2x, Kapa Biosystems). [N] cDNA PCR cycles were performed for each sample, where N was based on the Cycle threshold (Ct) value obtained from the diagnostic qPCR. After ssPCR, the samples were cleaned by adding 90 μl AMPure XP beads and washed three times according to the manufacturer's protocol. cDNA PCR products were eluted in 25 μl of water. The size distribution of the cDNA PCR products was checked with the Agilent BioAnalyzer and diluted to 70 pg/μl. For the subsequent tagmentation (slightly modified from (Poulain *et al*, 2017)), which simultaneously fragments and tags input DNA with adapter sequences, 1 μl of sample was mixed with 5 μl Tagment DNA Buffer (Illumina, US), 1 μl PEG, 2.5 μl Amplicon Tagment mix (Illumina, US) and 0.5 μl of PCR grade water. Samples were incubated at 55° C for 10 min and chilled to 10° C. After reaching 10° C, 2.5 μl of NT buffer was added to strip the enzyme of the cDNA and the sample was incubated at room temperature for 5 min. Next, the fragments were amplified using a limited-cycle PCR. Additionally, this PCR also added index sequences to the 3' end of the DNA fragments, thereby enabling the sequencing of pooled libraries on an Illumina Sequencing System. For this, 1 μl nanoCAGE custom S-series primer (10 μM) and 1 μl of a Nextera XT N-Series Index primer (N7xx, 10 μM), 7.5 μl Nextera PCR Mastermix and 3 μl water were added to each sample and amplified by PCR (72° C for 3 min, 95° C for 3 min , [95° C for 10 s, 55° C for 30 s, and 72° C for 1 min] x 15 cycles, 72° C for 5 min; CFX96 Touch, Bio-Rad, Sweden). The amplified libraries were cleaned with 22.5 μl AMPure XP magnetic beads (Beckman Coulter, Sweden) according to the manufacturer's instructions and cDNA was eluted in 20 μl nuclease free water. cDNA quantity was determined by Qubit and final library size distribution was checked with the High-sensitivity DNA kit on an Agilent Bioanalyzer. Prior to sequencing, library concentrations were adjusted to 10 nM and pooled together. Sequencing was done on HiSeq2500 (Illumina, US) in Rapid Run mode, producing 100bp single end reads.

MS-RanN6: 5'-TAGTCGAACTGAAGGTCTCCGAACCGCT CTTCCGATCTNNNNNN-3'

Template Switching Oligo: 5'-TAGTCGAACTGAAGGTCTC CAGCA[barcode][UMI]TATA(rG)(rG)(rG)-3'

MS-dir1F: 5'-TAGTCGAACTGAAGGTCTCCAGC-3'

MS-dir1R: 5'-TGACGTCGTCTAGTCGAACTGAAGGTCTCC GAACC-3'

nanoCAGE Custom S-series: 5'-AATGATACGGCGACCACC GAGATCTACACTAGTCGAACTGAAGG-3'

Nextera XT N-Series: 5'-CAAGCAGAAGACGGCATACGAGA T[index]GTCTCGTGGGCTCGG-3'

#### nanoCAGE data processing and analysis

nanoCAGE reads were extracted using TagDust (Lassmann, 2015) (version 2.31) and those mapping to ribosomal RNA were removed. Reads were trimmed to remove Nextera XT N-Series 3' adaptors using cutadapt version 1.9.1 (Martin, 2012) with the maximum read-adapter mapping error rate set to 0.15, the minimum overlap length set to 1 and trimming of up to 4 adaptors was allowed. Remaining reads shorter than 25 bases were excluded. Duplicated reads with identical UMI and sequence (in the first 25 bp) were assumed to be PCR duplicates and only one was kept for further analysis. A first alignment to the mouse genome mm10 was performed using Bowtie (version 0.12.7) (Langmead *et al*, 2009) allowing no more than 2 mismatches in the first 28 bp on the 5' end of the read. Reads which aligned uniquely (selected with parameters -m 1 –best –strata) and reads which did not align underwent a second alignment to a custom index of 5' UTR sequences. For this index, refSeq 5'UTR sequences (O'Leary *et al*, 2016) were extended with 78 bases upstream genomic and downstream mRNA sequences (78 is the maximum length of nanoCAGE reads after trimming of all adaptors). All best strata alignments with a maximum of 3 mismatches in the first 25 bases of the read were reported. One or two bases were trimmed from the 5' end of each read when alignment mismatches were present at the first or second position, respectively. Strand invasion artefacts were removed using the perl script provided in Tang et al. (Tang *et al*, 2013) with minimum edit distance set to 2.

Transcription start site peaks (defined as 5' end mapping positions of reads) observed in less than 4 out of 6 samples (3 replicates and 2 barcodes per replicate) were excluded. Reads from all replicates were then merged. RefSeqs with less than 10 reads were excluded. The complexity was verified by sampling increasing numbers of reads (delta of 150 000) and counting the number of peaks (>2 reads) and number of RefSeqs (>20 reads) detected. An average out of 3 iterations was used. For each gene entrez identifier, the RefSeq(s) with most reads was (were) selected (for multiple RefSeqs with the same maximum counts, an average was considered for each 5' UTR characteristics). The 5' UTR length was computed as a mean length along the RefSeq weighted by the read count of each observed peak. We then extracted the 5'UTR sequence corresponding to the most prevalent TSS position and computed GC content, fold energy and identified potential uORF with strong Kozak context (a start codon and in-frame stop codon within the 5' UTR; the start codon being in a [A,G]..ATGG context). Fold energies were obtained using mfold version 3.6 with default parameters (Zuker, 2003). We considered in the analysis herein the minimum folding energy after re-evaluation by the efn algorithm (see mfold manual). Ten UTRs failed to be in silico folded by mfold (these sequences were amongst the shortest ones) and positive fold energy results (n = 23) were excluded.

The weighted length, GC content and fold energy distribution of 5'UTR were compared between mRNAs regulated via different modes and visualized as boxplots. Wilcoxon-Mann-Whitney tests were used to assess differences between translationally offset and non-offset (mRNA abundance) mRNAs. Presence of strong uORFs is shown as a bar chart and differences tested using Fisher's exact tests. A global Bonferroni adjustment was applied on p-values.

#### Preprocessing and analysis of RNAseq of small RNAs data

The 3' adapter TGGAATTCTCGGGTGCCAAGG was trimmed when present using fastx_clipper (part of FASTX Toolkit version 0.0.14). Reads which were less than 17nt-long were excluded. Reads longer than 23 nt were also filtered out. Remaining reads were aligned to the mm10 mouse genome using Bowtie version 1.0.1 (Langmead *et al*, 2009) allowing one mismatch anywhere in the sequence. All alignments falling in the best alignment stratum were reported (parameters -v 1 –best –strata -a). miRNA levels were quantified using the countOverlaps function of the GenomicAlignments R/Bioconductor package (Lawrence *et al*, 2013) and the miRBase definition of miRNA coordinates Release 22 (Kozomara & Griffiths-Jones, 2014). For miRNAs mapping to multiple genomic locations, the quantification with minimum total count was considered. Only mature miRNA counts were kept in the analysis. miRNAs with average count below 30 were excluded. Differential expression analysis upon ERα depletion was performed using DESeq2 (Love *et al*, 2014) on raw miRNA counts without independent filtering. The replicate identifier was included as a covariate in the models. miRNAs were considered significantly differentially expressed for FDR<0.2 and abs($\log_2$ Fold Change) > $\log_2(1.25)$.

Datasets of targets of conserved mouse miRNAs were downloaded from TargetScanMouse release 7.1, January 2016 (Lewis *et al*, 2005) from tables "miR Family" and "Predicted Conserved Targets". Only highly conserved (code 2) and conserved (code 1) miRNA families were selected. Each mouse miRNA was then mapped to mRNAs having conserved target site(s) for this miRNA. Next, we tested whether downregulated miRNAs would have target sites overrepresentation among upregulated but translationally offset vs. upregulated non-offset mRNAs using Fisher exact tests. A Bonferroni correction was applied on Fisher exact p-values. To visualize this overrepresentation, each upregulated mRNA upon ERα depletion (mRNA abundance Up or Offset (mRNA Up)) was assigned an angle given by arc-tangent (polysome-associated mRNA log2 Fold Change / cytosolic mRNA log2 Fold Change). This corresponds to the angle between the horizontal reference line and the position of this mRNA in Fig. 1E (see Fig. 4E, blue arrow). For each miRNA, the distribution of its targets' angles is plotted together with the background (mRNAs non targets of any downregulated miRNAs). We proceeded with the same approach for upregulated miRNAs and downregulated mRNAs. We also compared the distributions of angles of mRNA targets of any upregulated miRNAs (10 miRNAs were upregulated) with mRNA targets of groups of 10 randomly selected miRNAs (we generated 30 such groups of 10).

#### Analysis of codon usage

For all mRNAs categorized under the different modes for regulation of gene expression, the longest coding sequence was retrieved from the consensus coding sequence project database release 21 for NCBI mouse annotation release 106 and Ensembl release 86 (2016-12-08, (Pruitt *et al*, 2009)). Once the average number of each codon across mRNAs under each mode for regulation of gene expression was computed; methionine, tryptophan and stop codons were removed and a Chi-Square test of independence was performed. The standardized residuals from this test were visualized using a heatmap. Rows and columns of the heatmap were ordered according to unsupervised hierarchical clustering using the 1 - Pearson correlation distance and average linkage. All p-values were adjusted for multiple testing using the Bonferroni correction and the significance threshold for being colored in the heatmap was 0.01. The top 12 codons with highest positive residuals (i.e. most over-represented codons in a mode of regulation) were selected for further analysis. This resulted in 9 codons over-represented among Offset (mRNA Up) and 3 codons over-represented among offset (mRNA Down).

Five measures of codon bias were performed at the mRNA level using the codonW software version 1.4.4: the Codon Adaptation Index (Sharp & Li, 1987), the Frequency of Optimal codons (Ikemura, 1981), the Codon Bias Index (Bennetzen & Hall, 1982), the effective number of codons (Wright, 1990) and the G+C content at the 3rd position of synonymous codons. The tAI was calculated using the tAI R package (dos Reis *et al*, 2003, 2004). The tAI measures the adaptation of the codon composition of the tRNA pool. Relative tRNA levels are estimated from copy numbers of each tRNA in the genome. tRNA genes from GRCm38/mm10 (Dec. 2011) were retrieved using the UCSC table browser and used as input (Karolchik *et al*, 2004). The tAI was also computed on coding sequences divided into sextiles. The distribution of these codon measures was compared between the different modes of regulation.

Codon composition of GO terms (Biological processes only) was plotted in the two first dimensions of a correspondence analysis using the FactoMineR R package (Lê *et al*, 2008). GO terms with less than 40 genes or more than 1000 genes were excluded and the R/Bioconductor package org.Mm.eg.db version 3.6.0 (Carlson, 2018) was queried to get go_id to entrez id associations (genes with IEA as only association were excluded as described above). Codon composition of each mode for regulation of gene expression was projected on this correspondence analysis. The FactoMineR package was also used to perform a gene-level correspondence analysis of codon composition of all regulated genes (from Fig. 1E).

When analyzing codon usage normalized by amino acid (each codon count was divided by its corresponding amino acid count), the data was scaled to account for the fact that amino acids are encoded by different numbers of synonymous codons.

In order to analyze which codons showed general co-usage in the same mRNAs, we used the Spearman rank correlation of each pair of codons within genes in all modes of regulation.

#### Quantification and analysis of tRNA levels

tRNA quantification was performed using the RNAseq of small RNAs using the ARM-seq bioinformatics pipeline from Cozen et al. (Cozen *et al*, 2015) with few modifications. Briefly, the tRNAscan-SE algorithm was used to predict the presence of tRNA loci in the mm10 genome. These tRNA loci were "in-silico processed" (introns are removed, CCA is added, 5-prime G base is added to Histidine tRNAs, N-bases are added to the ends of each mature sequence) using the maketrnadb script. All tRNA loci and mature tRNAs were added to the mm10 genome and a Bowtie2 index was created for alignment of the RNAseq reads of small RNAs after adaptor trimming (same trimming method as for the miRNA pre-processing). We used the default Bowtie2 alignment strategy (i.e. search for multiple alignments, report the best one, with upper limits for parameters –D and –R set at 100 (Langmead & Salzberg, 2012)). The quantification was performed using the ARM-seq countreads.py script. All reads mapping tRNAs were retained, regardless of where in the tRNA they mapped. DEseq2 was used for read count normalization (quantification of non coding RNA features retrieved from RNAcentral Release 10 (The RNAcentral Consortium, 2017) was provided to estimate size factors) and analysis of differential expression (Love *et al*, 2014). Mature tRNA raw counts were summed for each anticodon and the differential expression analysis was performed on all non coding RNA with a minimum average count of 30. tRNA coverage was computed using the ARM-seq pipeline script.

#### Analysis of public dataset for E2 dependent expression of ELP3, ALKBH8 and CTU2

CEL files were downloaded from the GSE35428 GEO repository (Wardell *et al*, 2012). Gene expression was normalized using Robust Multiarray Averaging and annotated with the custom probeset definition HGU133Plus2_Hs_ENTREZG (http://mbni.org/customcdf/21.0.0/entrezg.download/). Gene expression of *ELP3*, *ALKBH8* and *CTU2* was presented as boxplots and compared between conditions with Ethanol, E2, ICI-182,780 and E2+ICI-182,780 (10 replicates per condition). Wilcoxon-Mann-Whitney tests were performed for all 3 genes and a global Bonferroni adjustment was applied on p-values.

#### Western blot

Samples were collected from cells grown to 70% confluence and lysed in RIPA buffer (Sigma) containing 5 mM NaF, 1 mM $Na_3VO_4$ and 1X complete protease inhibitor cocktail (Roche). Cell lysates were quantitated using a detergent compatible protein assay kit (BioRad). Cell extracts were denatured and separated by SDS-PAGE. Blots were incubated with the following antibodies overnight at 4°C, on a platform shaker; Estrogen Receptor alpha (ERα, 1:2000, [6F11] Leica), DEK (1:1000 [5E3A6], Protein Tech), DEK (1:1000, [E1L3V], Cell signaling), ELP3 (1:1000, [D5H12], Cell signaling), MYC (1:1000, [Y69], abcam), β-Actin (1:100,000, [AC15], Abcam), JAG1 (1:2000, D4Y1R, Cell signaling), CHK1 (1:1000, #2360, Cell signaling), DCXR (1:1000, [15188-1-AP], Protein Tech), Androgen Receptor (AR, 1:10,000, [D6F11], Cell Signaling], GAPDH (1:100,000, [6C5], abcam), α-Tubulin (1:200,000, [DM1A], ThermoFisher), MYC-Tag (1:2000, [9B11], Cell signaling), HA-Tag (1:10,000, [12CA5], Sigma). Blots were then incubated for 1 hour with 5% skim milk and anti-rabbit or anti-mouse, horse radish peroxidase (HRP) conjugated secondary antibody HRP (Dako) amplification. Blots were incubated with Immobilon Western Chemiluminescent HRP Substrate (Merck) and Chemiluminescence was detected with amersham hyperfilm ECL (GE Healthcare) film. Antibodies have been validated by knockdown and/or overexpression of target proteins (except for commercially well validated antibodies e.g. β-Actin, GAPDH).

#### Analysis of codon usage in a public dataset of Tamoxifen-sensitive and resistant cell lines

Pre-processed RNAseq reads were downloaded from the GEO repository (Geter *et al*, 2017). We only considered data from heavy polysome mRNA (mRNA associated with > 4 ribosomes in this study) and total mRNA. Filtering of genes with zero counts, normalization (default settings) and analysis of changes in translational efficiency leading to altered protein levels was performed with anota2seq (Oertlin *et al*, 2018). The models were fitted on data from all samples (2 replicate of 3 conditions: tamoxifen sensitive, tamoxifen resistant, tamoxifen resistant with eIF4E silencing) and only the "Tamoxifen-resistant vs sensitive" contrast was considered for codon usage analysis. The frequency of AAA, CAA and GAA codons were calculated for the coding sequences of all human genes. These genes were categorized into deciles of such frequencies. Empirical Cumulative Distribution Function of the changes in translational efficiency between Tamoxifen resistant and sensitive cells was plotted for each decile. Wilcoxon-Mann-Whitney tests were performed to compare each AAA, CAA, GAA codon frequency decile to the first decile.
